# System-level high-amplitude co-fluctuations

**DOI:** 10.1101/2022.07.26.501262

**Authors:** Richard F. Betzel, Evgeny Chumin, Farnaz Zamani Esfahlani, Jacob Tanner, Joshua Faskowitz

## Abstract

Edge time series decompose interregional correlations (functional connectivity; FC) into their time-varying contributions. Previous studies have revealed that brief, high-amplitude, and globally-defined “events” contribute disproportionately to the time-averaged FC pattern. This whole-brain view prioritizes systems that occupy vast neocortical territory, possibly obscuring extremely high-amplitude co-fluctuations that are localized to smaller brain systems. Here, we investigate local events detected at the system level, assessing their independent contributions to global events and characterizing their repertoire during resting-state and movie-watching scans. We find that, as expected, global events are more likely to occur when large brain systems exhibit events. Next, we study the co-fluctuation patterns that coincide with system events–i.e. events detected locally based on the behavior of individual brain systems. We find that although each system exhibits a distinct co-fluctuation pattern that is dissimilar from those associated with global events, the patterns can nonetheless be grouped into two broad categories, corresponding to events that coincide with sensorimotor and attention systems and, separately, association systems. We then investigate system-level events during movie-watching, discovering that the timing of events in sensorimotor and attention systems decouple, yielding reductions in co-fluctuation amplitude. Next, we show that by associating each edge with its most similar system-averaged edge time series, we recover overlapping community structure, obviating the need for applying clustering algorithms to high-dimensional edge time series. Finally, we focus on cortical responses to system-level events in subcortical areas and the cerebellum. We show that these structures coincide with spatially distributed cortical co-fluctuations, centered on prefrontal and somatosensory systems. Collectively, the findings presented here help clarify the relative contributions of large and small systems to global events, as well as their independent behavior.

## INTRODUCTION

The human brain is fundamentally a complex network [1]. At the large-scale, white-matter fiber tracts constrain the dynamic interactions between brain regions, shaping their activity and inducing correlations–i.e. static functional connectivity (FC) [2–4]. The organization of this functional network is impermanent, fluctuating from one moment to the next [5–7]. In most applications, sliding windows are used to estimate patterns of time-varying FC (tvFC) [8]. Although used widely, the windowing method can induce artifactually smooth and temporally imprecise estimates of functional network architecture [8, 9].

Recently, we proposed a method for decomposing static FC into time-varying components [10–12]. This “edge-centric” approach obviates the need for sliding windows and returns temporally precise estimates of the instantaneous co-fluctuation between pairs of brain regions.

Previous applications of the edge-centric framework to resting-state fMRI BOLD data have reported “events”–short-lived and infrequent episodes in which many edges exhibit simultaneous, high-amplitude co-fluctuations [10]. Events, which have also been described using other methods [13–15], arise from the underlying modular structure of anatomical connectivity [16] and the co-fluctuation patterns they express are strongly correlated with static FC [10], carry subject-specific information [17], have been linked to endogeneous hormone fluctuations [11], and in some cases improve brain-behavior correlations [10].

However useful events may be, a narrow range of methods have been used to detect and characterize them. In particular, events have been defined *only* at the global scale. That is, for a frame to be classified as an event, the magnitude of co-fluctuations across the entire brain must exceed some statistical threshold [17]. Consequently, potentially neurobiologically meaningful co-fluctuations among smaller subsets of edges are unlikely to register as an event, even if their magnitude is exceptional strong.

Here, we seek to address this gap in knowledge by detecting events not at a global scale, but at the level of canonical brain systems [18, 19] and network modules [20]. We find that, as expected, the timing of global events is decoupled from that of small brain systems but correlated with co-fluctuations involving edges within the largest networks. We also find that events at the system level give rise to stereotypical topographic profiles and can be grouped into two broad clusters–one corresponding to events in sensorimotor and attentional systems and another by higher-order association systems. Next, we show that system level events can be used to generate partitions of edges, which can be used to extend system boundaries and allow for overlapping system-level architecture. These results add nuance to the growing literature on both edge-centric network analyses and neural events, while opening up avenues for future work.

## RESULTS

Here, we analyze resting-state data from the Human Connectome Dataset [21]. Specifically, we focus on a previously identified subset of 172 participants with low in-scanner motion [22] (of these, we analyze the 156 that had usable data for all scans). In addition, we also analyzed 7T movie-watching data, comparing it with restingstate data acquired at a similar field strength. We also cross-validate subsets of our findings using a third dense-sampling dataset: the Midnight Scan Club, comprised of ten individuals scanned ten times each [23, 24]. In all cases, we focus on a parcellation of the data in which the cerebral cortex, cerebellum, and subcortical nuclei were sub-divided into *N* = 482 parcels (*N_cortex_* = 400; *N_cb_* = 32, *N_subcortex_* = 50) [25–27]. Across all datasets, we estimated edge time series (Fig. 1a-d) and, using a data-driven procedure, detected events globally as peaks of the whole-brain co-fluctuation amplitude (root mean square – RMS; Fig. 1e,f). We also used the same algorithm to detect local events for 24 putative brain systems (16 cortical; 1 cerebellar; 7 subcortical; see Fig. S1 for surface and volumetric definitions of systems) by restricting the algorithm to use only edges that fall within brain systems (Fig. 1g).

**FIG. 1.**
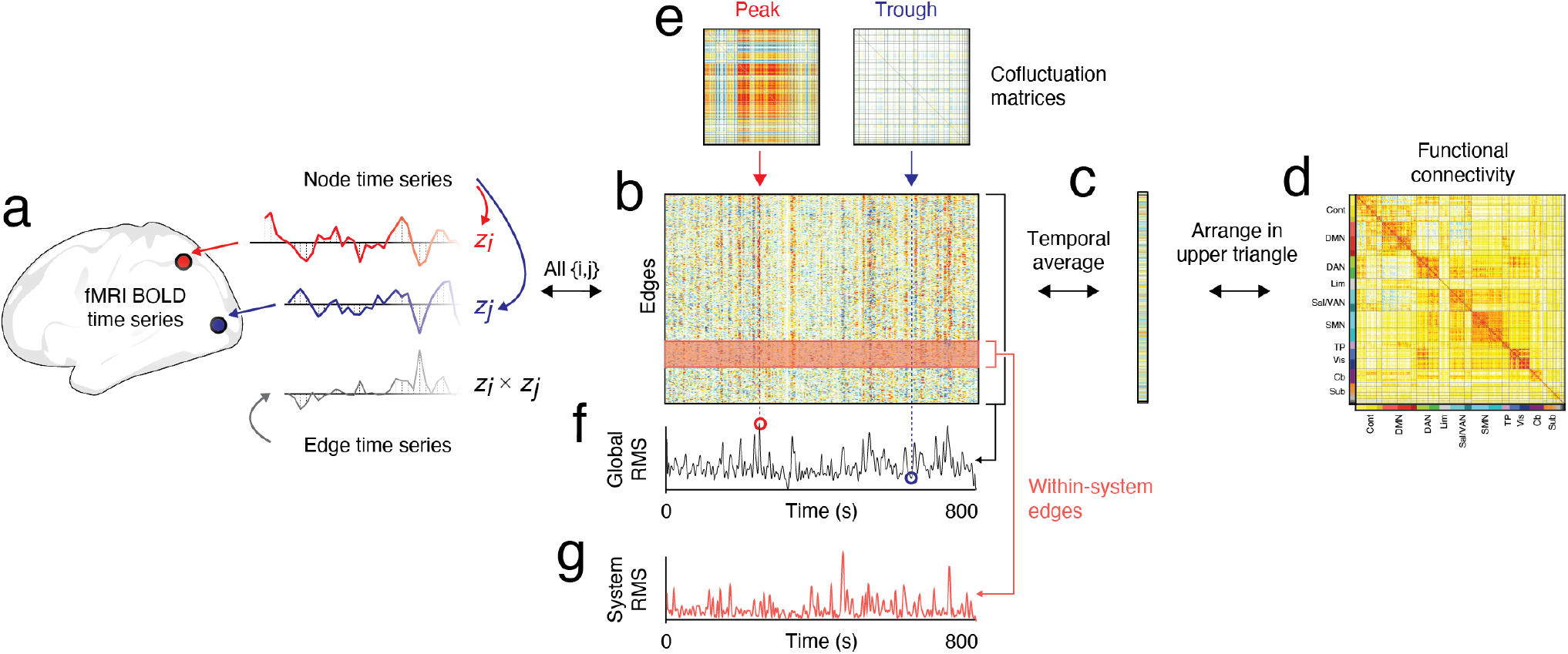
Schematic of edge time series and global/system co-fluctuation amplitude calculations. (*a*) Edge time series are calculated as the element-wise product of z-scored regional time series. (*b*) This procedure can be repeated for all edges. (*c*) The temporal average of an edge time series is a correlation; averaging the edge time series for all edges yields an exact estimate of the correlations for all edges (in vectorized form). (*d*) The elements of this vector can be rearranged into the upper triangle of a region × region functional connectivity matrix. (*e*) A “slice” across all edge time series at time *t* yields an estimate of the instantaneous co-fluctuation pattern, which can be modeled as a network. There is variation in the overall magnitude of co-fluctuations from one time point to another. (*f*) The whole-brain co-fluctuation amplitude can be summarized as the square root of the mean of squared co-fluctuations (RMS). (*g*) An analogous RMS time series can be calculated for brain systems/modules by calculating the mean across only those edges that fall within a given system.

### Global events are driven by larger brain systems

Previous studies have defined global events – intermittent, brief, and high-amplitude co-fluctuation patterns – as peaks in the RMS time series whose amplitude is statistically greater than that of a null distribution [10, 11, 17, 28, 29]. However, global events may inadvertently be driven by the behavior of larger brain systems, whose collective amplitude contributes disproportionately to the whole-brain RMS signal. Here, we verify that this is the case, showing that, as expected, whole-brain co-fluctuations are strongly correlated with those of the largest systems.

We compared whole-brain and system-level RMS time courses. We found marked heterogeneity across systems in terms of how strongly their respective RMS was correlated with the global RMS (Fig. 2a). Importantly, the magnitude of correlation was closely related to system size (number of nodes); in general, larger systems were more strongly correlated (Spearman rank correlation; *ρ* = 0.84; *p* < 10^−15^; Fig. 2b). Notably, this effect was also evident when the RMS time series were binarized (events = 1, non-events = 0). Again, we found that the timing of locally-defined events – based on systems’ internal RMS compared to a null model – with global events was biased so that the correspondence was greatest for larger brain systems (*ρ* = 0.84, *p* < 10^−15^; Fig. 2c,d). Notably, these effects hold irrespective of whether global signal regression is included in the processing pipeline (Fig. S2a,b). For completeness, we also show the inter-system correlation matrix for both event time series (binary vectors of peak co-fluctuations) and RMS time series (Fig. 2e).

**FIG. 2.**
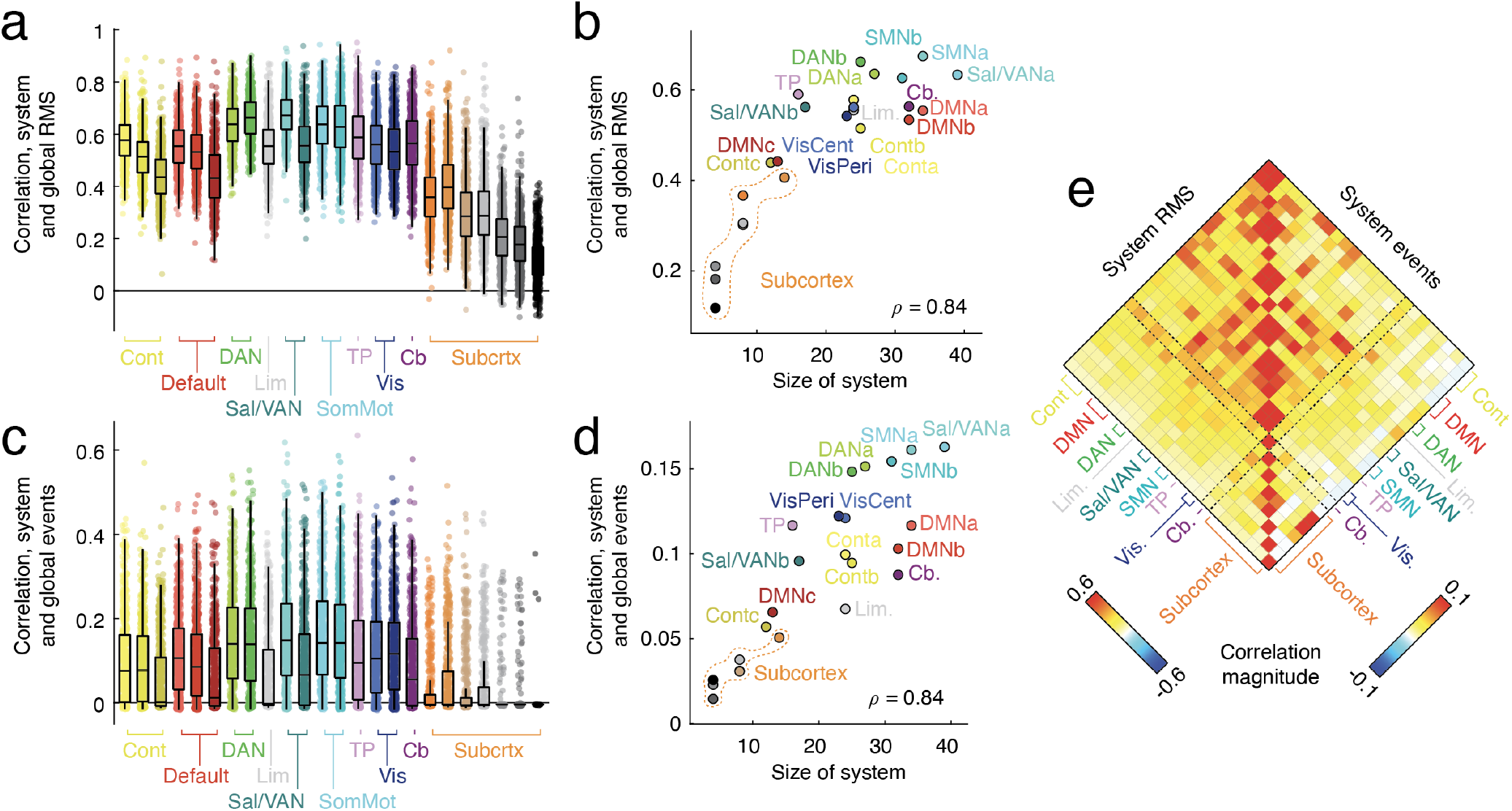
Correlations between system and global co-fluctuations. (*a*) Correlation magnitude of system-level and global RMS time series. Each point represents a scan of an individual. (*b*) Scatterplot of mean correlation between system/global RMS and system size. (*c*) Correlation between system and global binary event time series (all frames are 0 except for event peaks, which are assigned a value of 1). (*d*) Scatterplot of mean correlation between system/global event time series and system size. (*e*) Inter-system correlation structure of RMS (*left*) and event time series (*right*).

Together, these results suggest that global events – the focus of numerous studies to date – are largely driven by the time-varying co-fluctuations of the largest brain systems. This effectively renders “event detection methods” blind to the behavior of smaller, more focal brain systems, motivating the exploration of events at the local (system-level) scale.

### Statistics of system-level events

Results from the previous section suggest that, although all systems exhibit events, the amplitude and frequency with which they occur depends on system size. In general, larger systems comprised of more nodes tend to exhibit RMS co-fluctuations and events that are coincident with those defined globally. Here, we investigate properties of system-level events in greater detail.

Here, we analyzed system-level RMS data along with the binarized event vectors described in the previous section (Fig. 3a,b). We found that sub-cortical systems exhibited greater RMS during events compared to cortical systems (*t*-test; *p* < 10^−15^; Fig. 3c). This effect was reversed when we calculated the event rate for each system, i.e. the likelihood that a frame would be classified as an event based on the system’s co-fluctuations, with cortex exhibiting a much greater event rate than subcortex (*t*-test; *p* < 10^−15^; Fig. 3d).

**FIG. 3.**
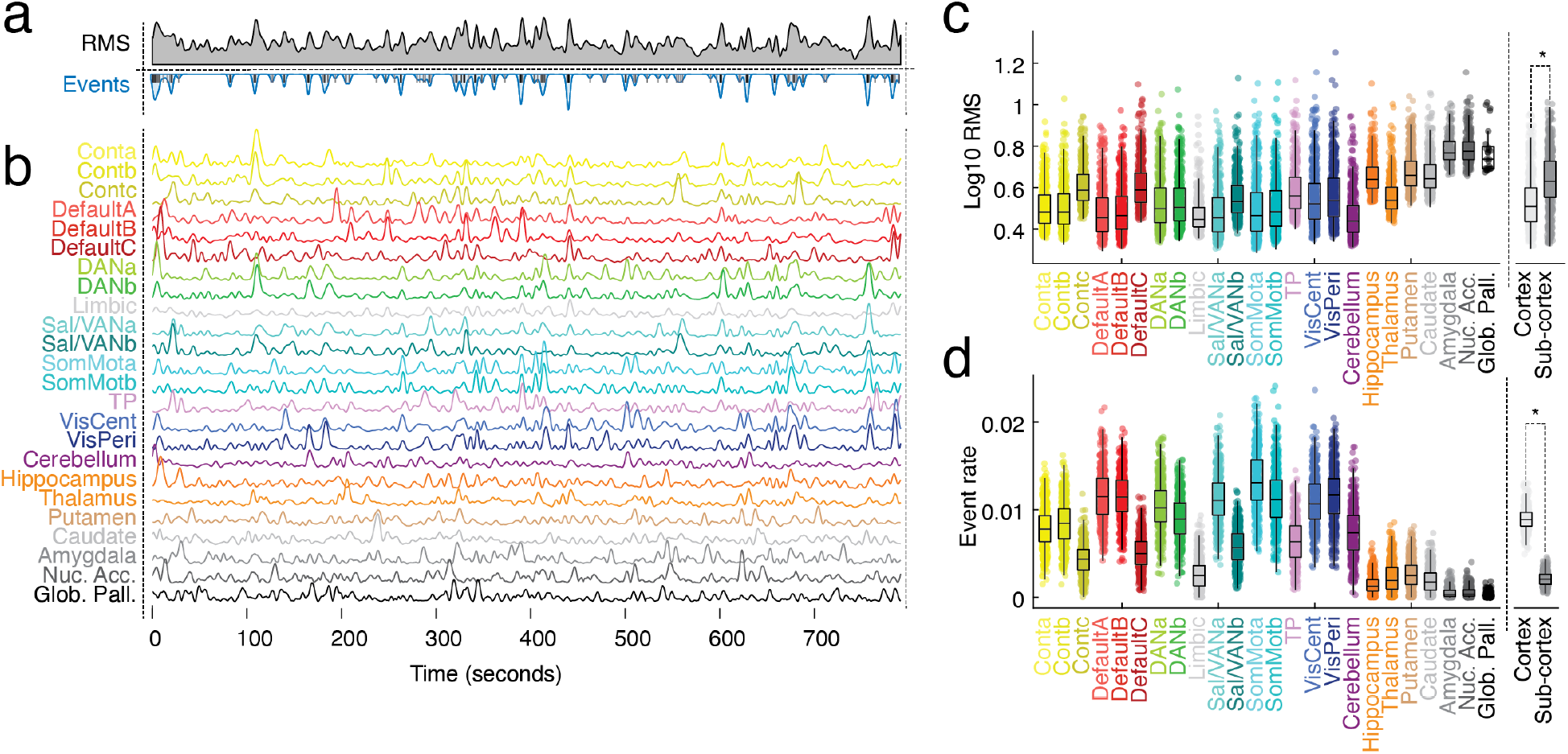
Statistical properties of system-level co-fluctuations. (*a*) Example global RMS (*top*); summed and, for visualization only, smoothed system-level event time series (*bottom*). (*b*) Example system-level RMS time series. (*c*) RMS amplitude of peak co-fluctuations (those classified as events). Each point represents an individual event from a particular subject/scan. While the median, interquartile range, and whiskers were calculated based on the full distribution, we restricted the number of points displayed in each plot for the ease of visualization. (*d*). Event rate (fraction of frames classified as events) for each system.

What explains the variation in event rates and peak RMS across systems? One looming factor is a system’s size and its interaction with the null model used to detect events. Here, we compared observed RMS values against a null distribution generated by independently circularly-shifting each region’s time series, estimating edge time series from the shifted data, and calculating its RMS. This procedure was repeated, yielding a null distribution. In general, as system size increases, the sampling error around the mean RMS narrows, i.e. with very large systems the variability of system-averaged RMS gets smaller. While the criterion for statistical significance does not vary with system size, the effective RMS cutoff does, with smaller systems requiring greater RMS values to classified as an event (see Fig. S3a). Other factors that influence peak RMS and event rate are related to the time-invariant correlation structure of nodes in a given system. Systems with greater mean FC exhibited lower peak RMS (*ρ* = −0.21; Fig. S3b) but higher event rates (*ρ* = 0.81; Fig. S3d), while systems whose interregional correlations were more variable exhibited reduced peak RMS (*ρ* = −0.52; Fig. S3c) and greater event rates (*ρ* = 0.61; Fig. S3e).

Collectively, these findings suggest that co-fluctuations of edges within cortical systems exhibit distinct co-fluctuation profiles but that care must be taken to not conflate effects of system size and its static FC properties with those specific to events.

### Whole-brain co-fluctuation patterns during system events

To this point, we have considered brain systems in isolation, focusing on the collective co-fluctuations of within-system edges. However, brain systems are coupled to one another and interact across time. When one system has an event, what other edges and systems “tag along” and also exhibit strong co-fluctuations?

To address this question, we examined “system event snapshots”. That is, given an event in system *s* at time *t*, we extracted the whole-brain co-fluctuation pattern at that instant (Fig. 4a). We subsequently averaged these patterns across individuals, scans, and instances, yielding average co-fluctuation maps (Fig. 4b; see Fig. S4 for projections of co-fluctuation maps onto cortical surface).

**FIG. 4.**
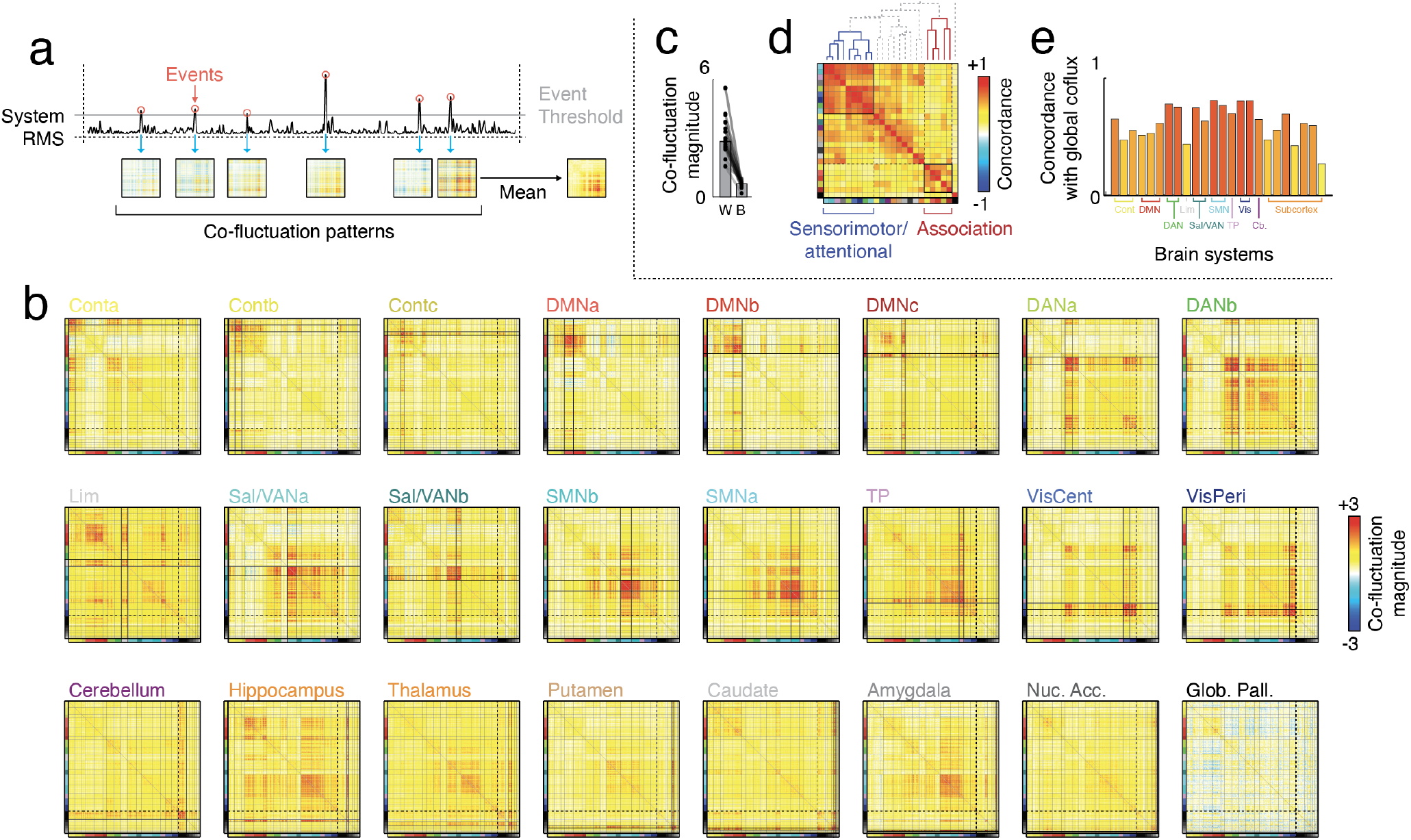
System event snapshots events and co-fluctuation matrices. (*a*) System events are detected by comparing the observed system RMS time series again a null distribution. Events are defined at the peak RMS of supra-threshold frames. (*b*) Average co-fluctuation matrices during system event snapshots. (*c*) Co-fluctuation amplitude for edges that fall within the system in which the event originated compared with all other edges. (*d*) Concordance matrix calculating the similarity between all pairs of snapshot co-fluctuation matrices. Dendrogram above the matrix indicates hierarchical relationships between pairs fof systems. (*e*) Similarity of each snapshot’s co-fluctuation matrix to the global co-fluctuation matrix.

As expected, the co-fluctuation magnitude was strongest between regions within the system in which the event originated (paired sample *t*-test, *p* < 10^−15^; Fig. 4c). However, there were also examples where events originating in system *s* exhibited non-local co-fluctuations, e.g. those involving other systems. For example, dorsal attention system (DANa) events coincided with strong co-fluctuations among central and peripheral visual systems. System event snapshots also exhibited high levels of specificity. Consider dorsal attention systems a and b (DANa and DANb, respectively). Although topographically interdigitated, DANa events occur coincidentally with visual events whereas DANb events coincide with co-fluctuations among regions in the somatomotor and salience/ventral attention networks (Fig. 4b).

Indeed, we find broad evidence that system events exhibit rich topographic profiles. To highlight this, we clustered co-fluctuation patterns hierarchically based on their spatial concordance with respect to one another (Fig. 4d). Although the hierarchical tree can, in principle, be cut at any level, we found that two specific clusters of snapshots persisted across multiple levels. The first was comprised of somatomotor, visual, and attention systems. Although the co-fluctuation patterns vary subtly and systematically, events originating in any of the systems comprising this cluster exhibited similar co-fluctuation patterns. The second cluster included control and default mode systems exclusively.

Finally, we compared event snapshots to the co-fluctuation pattern obtained during global, whole-brain events. As expected, we found that systems in the sensorimotor/attentional and association clusters behaved dissimilarly. Specifically, we found that the sensorimotor/attentional systems exhibited co-fluctuation patterns that were more similar to the global events compared to the association cluster (*t*-test; *p* = 5.2 × 10^−6^; Fig. 4e).

Importantly, we found that system event snapshots exhibited similar spatial patterning, irrespective of whether the global signal was regressed out of regional time series (Fig. S2c and Fig. S5b,c), across datasets (Fig. S6), and when the systems were defined based on subject-specific variations of the canonical set (Fig. S7) or using data-driven approaches, e.g. modularity maximization (Fig. S8). We also found that the specificity of system event co-fluctuations could be improved by subtracting out the co-fluctuation pattern expressed during global events (Fig. S5a) and that, in general, system events exhibit a high degree of individualization (Fig. S9). Lastly. we found no evidence supporting the hypothesis that system events follow distinct temporal trajectories or exhibit precise timing, i.e. an event in system *r* consistently precedes an event in system *s* and so on (Fig. S10).

### Revealing overlapping system-level architecture

In the previous section, we calculated the mean co-fluctuation patterns during system events. These patterns can be further leveraged to investigate the brain’s overlapping system-level architecture (see Fig. 5 for a schematic illustration). To do this, we performed the following set of analyses. We first calculated the mean system event co-fluctuation pattern for each cortical system, subcortical area, and cerebellum. Next, for each edge, {*i*, *j*}, we asked during which of the 24 co-fluctuation matrices was the co-fluctuation magnitude the greatest, retaining both the magnitude of that co-fluctuation (Fig. 6a) and the “winning” system (Fig. 6b). Repeating this procedure for every edge returned a system assignment for each *edge* in the network. As in [30], this set of edge-level clusters allows us to calculate, from the perspective of brain regions, overlapping nodal community assignments. That is, because a region’s edges can be assigned to many different clusters, that region can be viewed as having overlapping clusters. Unlike [30], cluster assignments are obtained using a “winner take all” heuristic, rather than a semi-supervised clustering algorithm. Here, we also examine whole-brain clusters that include subcortical and cerebellar regions.

**FIG. 5.**
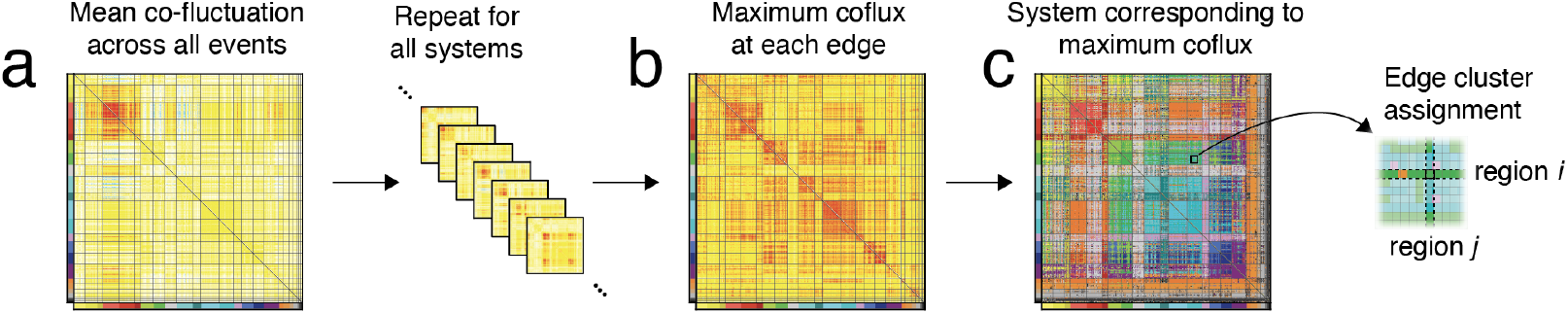
Estimating edge-level system assignments. (*a*) First, calculate the mean system event snapshot for all brain systems. Here, we show the mean pattern for DMNa.Repeating this procedure for each system yields 24 co-fluctuation matrices. For every edge, identify its maximum co-fluctuation magnitude across all systems (panel *b*) and the corresponding system (panel *c*). This yields a system assignment for each edge in the network. The inset shows labels for a subset of edges in greater detail.

**FIG. 6.**
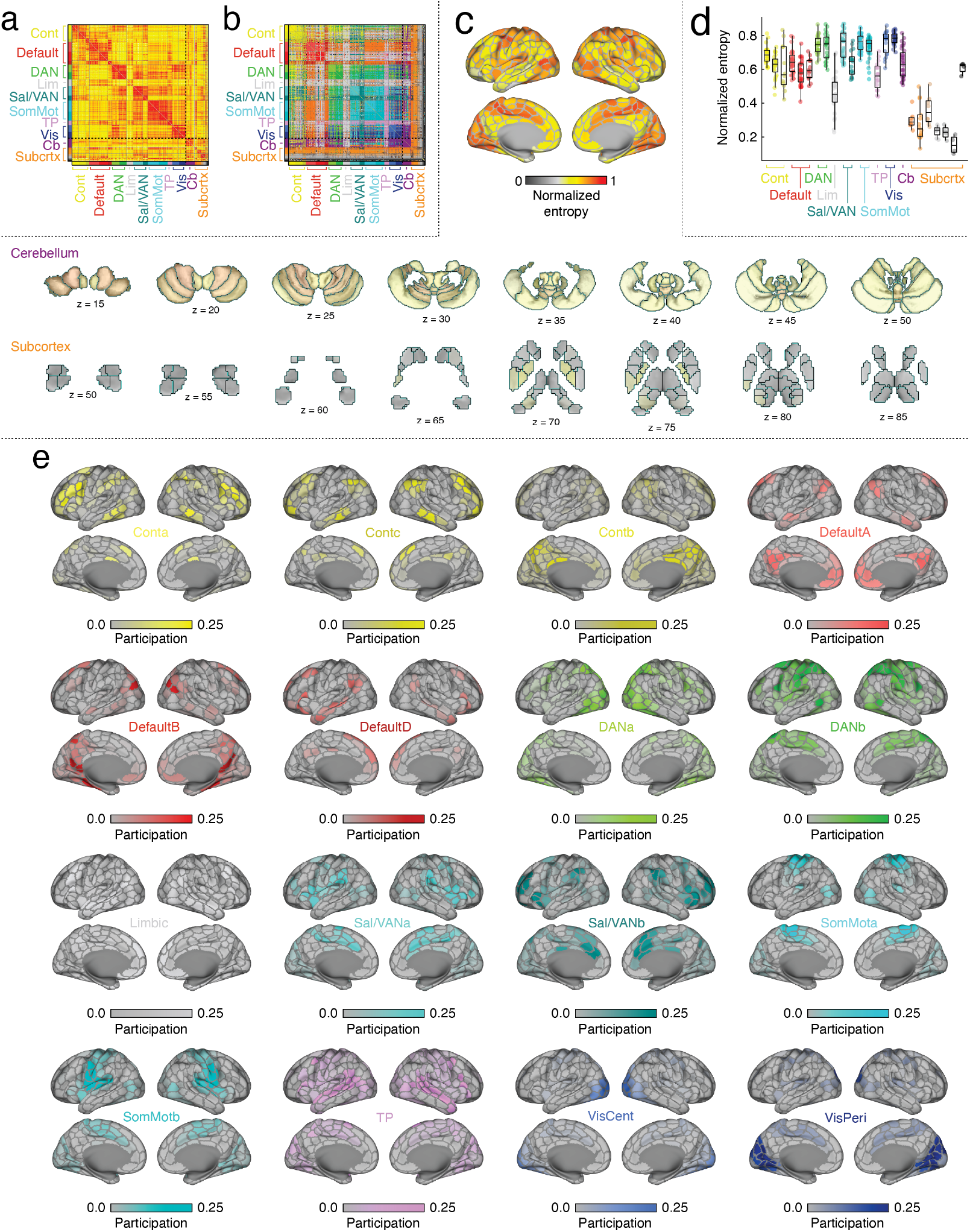
Overlapping system-level communities. (*a*) We calculated the maximum co-fluctuation magnitude for each edge across all system event snapshots. (*b*) We assigned each edge to the system whose events evoked its strongest co-fluctuation. (*c*) Given these edge-level system labels, we calculated each region’s overlap as an entropy – the extent to which its edges labels deviated from a uniform distribution. Here, we show entropy values projected onto cerebral cortex (*top*), cerebellum (*middle*), and subcortical regions (*bottom*). (*d*) Entropy values aggregated by brain systems. (*e*) Regional affiliation for each of the sixteen cortical systems.

Once we obtain cluster assignments, we can calculate each region’s overlap as a normalized entropy measure. Intuitively, regions whose edges’ cluster assignments are uniformly distributed have entropy values closer to 1. Regions whose edge cluster assignments all have the same value have entropy values closer to 0. Here, we find that entropy is heterogeneous across the brain, with the greatest levels of overlap concentrated in sensorimotor and attentional systems (Fig. 6c). Compared to cerebral cortex, we find relatively lower levels of overlap in subcortical systems, although the cerebellum exhibits comparable entropy levels (Fig. 6c,d). We show example overlap patterns – the fractional association of nodes with given systems – in Fig. 6e.

An important concern is that the overlap scores could be driven by weak co-fluctuations. That is, edges whose co-fluctuation pattern is uniformly weak across all system events and therefore have no clear system affiliation. To address this concern, we systematically removed weak co-fluctuations and recalculated the regional entropy scores using only the surviving edge cluster assignments (Fig. S11a). In general, we found that the entropy map was preserved over a wide range of thresholds (Fig. S11b,c). Another concern is that the entropy maps are driven by the behavior of within-system edges. Indeed, these edges were used to define system events and are almost always assigned to their original cluster (of the 5945 within-system edges 5705 had assignments to their original system; 96.0%). To avoid circularity in our analyses and to reduce the impact of these within-system edges on entropy estimates, we performed three analyses. First, we recalculated entropy values but excluded all within-system edges and found a nearly identical entropy map (Fig. S12). Complementing this analysis, we next calculated the extent to which each system’s overlap pattern was driven by within-system edges (Fig. S13), discovering heterogeneity across systems, but with the greatest off-diagonal contributions associated with hippocampal and limbic systems. Finally, we went one step further, and fully recalculated the “winning” system for each edge after censoring in each system event’s co-fluctuation matrix, all of the edges where at least one stub node is associated with the corresponding system (Fig. S14). Despite an entirely new set of edge system labels, we found that results were largely consistent with those reported earlier in this section using edge system labels estimated from the “intact” co-fluctuation matrices.

Collectively, we posit a novel method for assigning system labels to edges that reduces computational burden associated with existing techniques. Our findings suggest that sensorimotor and attentional systems exhibit the most varied and diverse edge-level assignments, an effect that holds across thresholds, when system edges are ignored, and when systems are defined using subject-specific variants of a group-level atlas and data-driven clustering algorithms.

### System-level events during movie-watching

To this point, we have only considered system-level events at rest – i.e. in the absence of stimulation or explicit task instruction. How do brain systems and the timing of their events vary during other experimental paradigms? Here, we begin to address this question by examining system-level events during passive movie-watching. Here, we analyzed movie-watching data from the Human Connectome Project (*N* = 172 subjects). In each scan, participants watched a series of concatenated movie scenes. Between every pair of scenes was a short rest period. In a recent study [29], we focused on global, whole-brain events and demonstrated that, although they occurred throughout the scan, there was a strong tendency for these events to occur during the rest periods. However, it is unclear how the timing of local (system-level) events coincides with the global events.

We analyzed these data exactly as before, calculating global and system-level co-fluctuation time series to identify the timing of putative events (Fig. 7a,b). We found that, compared to rest, movie-watching exhibited significant intersubject correlations in terms of system-level RMS time courses (Fig. S15). Movie-watching also prompted decoupling of dorsal attention, temporo-parietal, and visual systems from one another (Fig. 7c). As a consequence of this decoupling, events originating in any of these systems exhibited significantly lower co-fluctuation amplitude during movie-watching compared to rest (Fig. 7). Essentially, fewer systems participated in each event.

**FIG. 7.**
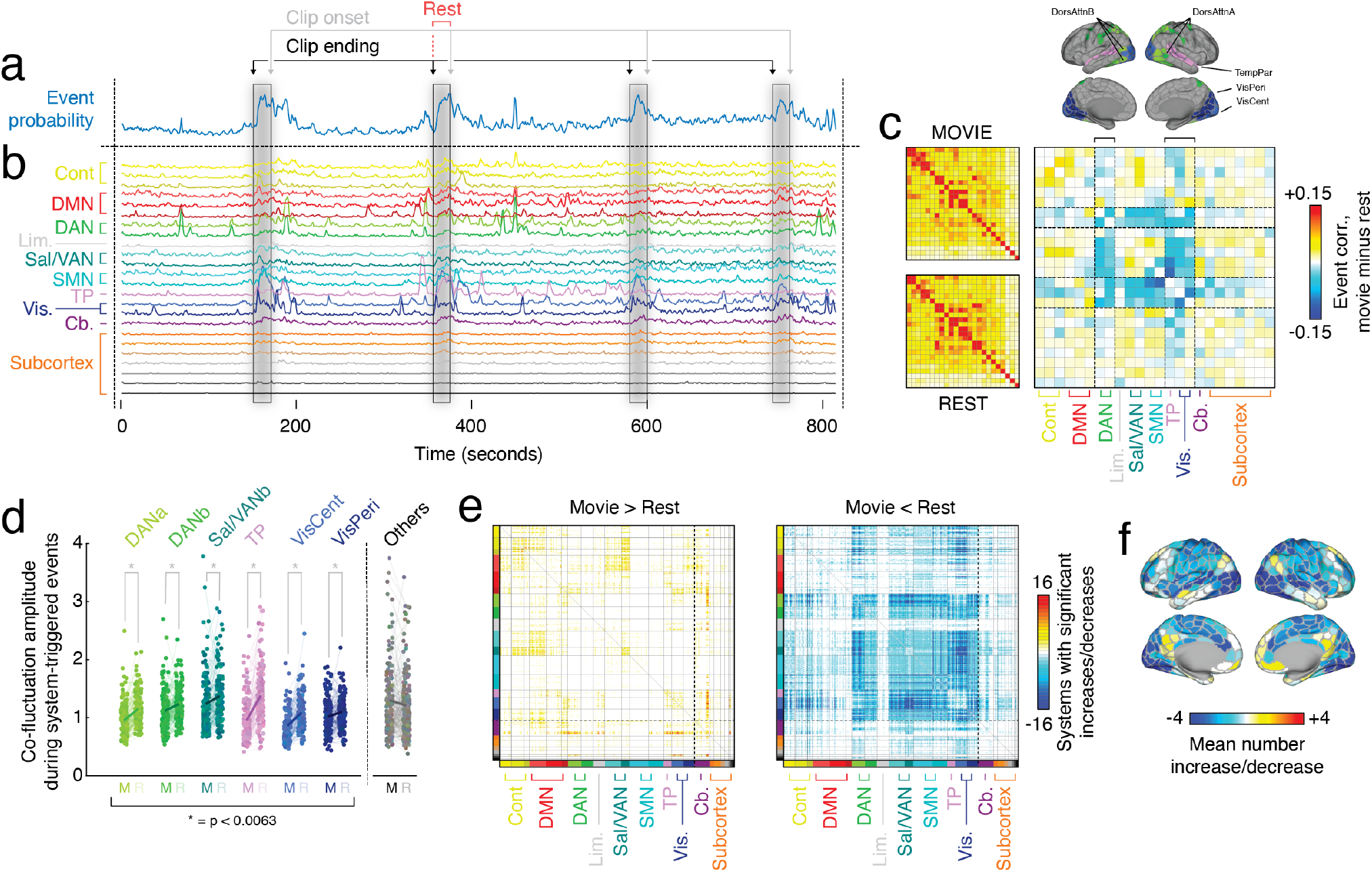
System-level events during movie-watching. Schematic illustrating timing of global, whole-brain events (panel *a*) and system-level events (*b*) during movie-watching. Note that, in general, global events are most frequent during rest periods between movie clips, whereas system-level events occur during rest but also during other points within each clip. (*c*) Correlation structure of the timing of system-level events during movie-watching (*top*) and during resting-state scans (*bottom*). We also show the difference in correlation structure (movie minus rest; *right*). The differences show that the timing of events in dorsal attention, temporoparietal, and visual systems decouple from one another during movie-watching, i.e. become more independent. (*d*) Co-fluctuation amplitude of system events during movie-watching and rest (denoted by ‘M’ and ‘R’). (*e*) Region pairs whose co-fluctuation magnitude increases/decreases during movie watching (interquartile range of movie minus rest across subjects excludes a value of zero). (*f*) Row mean of the matrices depicted in panel e and projected onto cortical surface.

When we examined the system events in greater detail and compared edge weights between rest and movie-watching, we found very few significant increases in co-fluctuation magnitude (movie over rest), but widespread decreases (Fig. 7e). The edges that increased and decreased were largely restricted to neocortex (but note the strong increases between cerebellar regions and a select set of cortical areas; Fig. 7e, left). Indeed, the decreased co-fluctuation changes were concentrated in a subset of the same brain systems that decoupled from one another, namely dorsal attention (DANa) and visual systems (Vis-Cent) (spin test, false positive rate fixed at *q* = 0.05; *p_adj_* = 0.014; Fig. 7f). Interestingly, because the decoupling and reduced co-fluctuation magnitude was *so* pervasive across the brain, the systems that increased (even modestly) all survived statistical testing. These include all sub-components of the default mode (DMNa,b,c) and a sub-component of the control network (Contb).

Taken together, these findings suggest that system-level events reconfigure their timing during movie-watching, decoupling systems for attending and processing exogenous stimuli from one another.

### Cortical responses to sub-cortical events

In the previous sections, we examined system-level events at the whole-brain level, pooling data from cortical, cerebellar, and subcortical sources. However, the interplay between cortical and non-cortical areas is complex, with many studies reporting area-specific static FC [31–33]. Here, we focus on the event structure of the cerebellum and basal ganglia, emphasizing the cortical responses to events originating in non-cortical areas.

To do so, we first calculated the global cortical co-fluctuation during events originating in subcortical and cerebellar systems (Fig. 8a). We found that, of all non-cortical systems, hippocampal events evoked the strongest cortical response. Evidence supporting this claim was apparent at the group level (Fig. 8b), but also at the subject level after accounting for inter-individual differences in mean co-fluctuation amplitude (Fig. 8c). Next, we calculated which of the non-cortical systems evoked the strongest co-fluctuation at each edge. Note that this approach is analogous to what was reported in Fig. 6b, the distinction here is that we excluded cortical systems. Again, we found that hippocampus, followed by thalamus, caudate, and cerebellum, evoked the strongest co-fluctuations in the greatest number of edges (Fig. 8d).

**FIG. 8.**
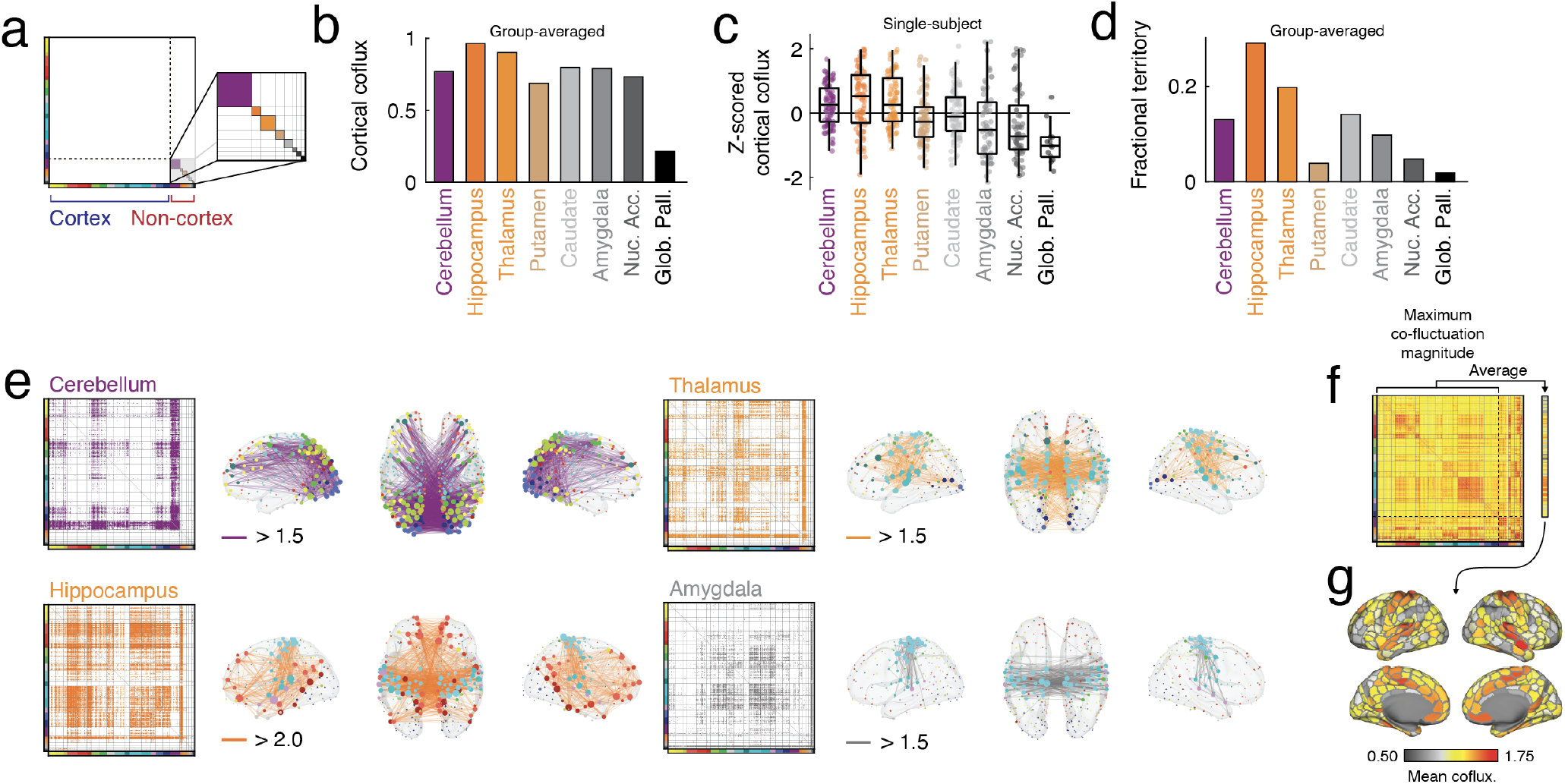
Cortical responses to non-cortical events. (*a*) Schematic illustrating the division of the connectivity matrix into cortical and non-cortical nodes. The inset depicts the eight non-cortical systems and their divisions in greater detail. (*b*) Mean cortical co-fluctuation amplitude in response to non-cortical events using group-averaged data. (*c*) An analogous plot but at the level of single subjects. Each point represents a subject. Note that prior to plotting, subjects’ evoked co-fluctuation magnitudes were z-scored (within subjects) to remove any subject level differences in mean co-fluctuation. (*d*) For each edge, we identified the non-cortical system whose events triggered the greatest cortical co-fluctuation amplitude. We show the fraction of edges associated with each of the eight non-cortical systems. (*e*) For a select set of non-cortical systems, we show the cortical edges that respond to their events. Here, we show edges with mean co-fluctuation above a particular threshold whose value varies from system and is shown in each sub-panel. Node size is proportional to the mean thresholded co-fluctuation magnitude of each node. (*f*) The maximum co-fluctuation for each edge across all non-cortical system event. (*g*) The mean regional responsivity - average of rows (or equivalently, columns) of the matrix shown in panel *f*.

Next, we asked which systems/regions exhibited the strongest evoked responses coincident with non-cortical events. To address this question, we calculated the maximum co-fluctuation of edges across all non-cortical event categories (Fig. 8d). Visually, we find that the strongest responses were concentrated in somatomotor and default mode systems and the edges that fall between them. We confirmed this by calculating the mean co-fluctuation (the row average) across all columns, and found that, as expected, the regions whose cortical responses to non-cortical events were strongest were concentrated in the aforementioned systems (Fig. 8e). Analogous to the analysis and data presented in Fig. 6a, we calculated the maximum co-fluctuation magnitude for edges during non-cortical events (Fig 8 f). From this matrix, we averaged the rows to ascertain the mean reactivity for each node, and plotted this on the cortical surface (Fig. 8g).

## DISCUSSION

Previous studies used edge time series to “temporally unwrap” functional connections, revealing bursty behavior across time and global events. The focus on whole-brain events has come at the expense of focal (system-level) high-amplitude co-fluctuations, which (for small systems) are unlikely to register as an event using existing event detection methods. Here, we address this limitation and focus, explicitly, on the co-fluctuation patterns of brain systems defined canonically and using data-driven techniques.

### Global events coincide with high-amplitude co-fluctuations in large brain systems

We found that as a system’s size increased, the amplitude of its co-fluctuations and timing of its events become strongly correlated with the global RMS signal and the timing of whole-brain events. This observation has two immediate implications. First, it suggests that the global events characterized in previous studies are likely driven by the co-fluctuations of large brain systems [11, 17, 28, 29]. Indeed, the event co-fluctuation patterns described in those studies are remarkably consistent in that they implicate large and distributed brain systems, including the default mode, control, and cingulo-opercular networks (or territories that are typically associated with those systems; see [34, 35] for discussions on naming and describing system-level architecture). Note also that the contribution to the global RMS from any given system grows supralinearly as a function of system size (approximately as the square of the system’s size). This is because a system of *N_s_* nodes contains 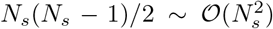 within-system edges. Consequently, large systems make disproportionately large contributions to the global RMS.

The second implication is that, if global events are driven by large brain systems, then the behavior of smaller systems has been under-reported in the extant literature. Indeed, we find evidence that, although broadly similar, events originating in individual systems are heterogeneous and reflect distinct coupling patterns. This opens up opportunities for future studies to explore the fine-scale temporal variation of small systems in future studies.

### System events are variable and exhibit hierarchical structure

Here, we examined system event snapshots–the whole-brain co-fluctuation patterns expressed whenever an event is detected within a given system. We made a series of observations. First, we found that co-fluctuation patterns were dominated by within-system effects. When system s exhibits an event, the strongest co-fluctuations (as expected) are among edges within that system. This effect is a mathematical necessity, as it is precisely the strong co-fluctuations among this ensemble of edges that defines an instant in time as an event.

Second, we noted that there were also strong co-fluctuations involving between-system edges. In other words, when system *s* exhibits an event, there is a tendency for other systems to simultaneously co-fluctuate. It is precisely these between-system co-fluctuations that cause static FC to deviate from a perfectly modular structure, inducing cross-system correlations when averaged across time [36, 37]. Interestingly, we find that system event co-fluctuation patterns can be organized into two larger clusters based on their spatial similarity; these groups broadly recapitulate a bipartition of the brain into task-positive/-negative systems [38–40].

Third, we find no evidence, at the population level, that systems exhibit distinct staging or timing leading up to an event. This is in contrast with a number of studies that have reported consistent lag structure in spontaneous fMRI BOLD activity [41], functional connections [42], and motifs of brain-wide connectivity patterns [43]. One possible explanation for these diverging results is that system-level events examine the collective behavior across ensembles composed of hundreds of edges. Although this behavior is generally cohesive and correlated, the RMS across these ensembles necessarily washes over inter-edge variation, possibly obscuring temporal delays. A second possible explanation is that delays exist at the level of regional activity, but that when combined multiplicatively (as in the generation of edge time series), these delays are effectively destroyed.

### Interactions between cortical and non-cortical systems

To date, most studies of edge time series have focused on cerebral cortex [9, 10, 12, 16, 17, 28, 29, 44, 45]. Consequently, much less is known about community overlap and event timing in cerebellar and subcortical systems. There are, however, two notable exceptions. In [46], the authors clustered edge time series to reveal distinct edge communities that bridged linked subcortical and cortical areas to one another, interpreting these communities as channels of communication. In particular, the authors discovered that hippocampus and amygdala exhibited were coupled to cortex *via* one pattern of edge communities while striatal and thalamic areas were coupled *via* a dissimilar pattern. In [47], the authors use similar methodology to characterize striatial edge communities, reporting a decoupling of edge community structure from the static FC, suggesting that the temporal variation captured by edge time series may be driven by inter-areal differences in anatomical inputs from cortex.

Here, we contribute to this growing body of literature. First, we show that, compared to cerebral cortex, subcortical systems exhibit greater RMS during events but dramatically reduced event frequency. These differences can likely be attributed to attenuated signal-to-noise in inferior, non-cortical areas. In short, subcortical systems require a proportionally greater RMS level to be considered an event due to a broader distribution of background noise (greater variance).

However, we also showed that the timing of subcortical and cerebellar events is coincident with events among specific subsets of cortical systems. Of particular interest is the pattern of evoked cortical co-fluctuations to hippocampal events, which includes default mode and somatomotor systems. This pattern is in line with a recent study in which activation of the default mode was tightly coupled to “bursty” hippocampal activity and eventually linked to spontaneous replay, hinting that a similar mechanism could be at play here [48].

We also found evidence of preferential engagement of cortical systems by events originating in non-cortical systems. For instance, and as noted before, hippocampal events coincide with strong co-fluctuations between default mode and somatomotor systems. Similar phenomena are evident during events originating in other non-cortical systems; thalamic events induce co-fluctuations between somatomotor, salience/ventral attention, and visual regions; amydalar events induce co-fluctuations involving default mode and somatomotor systems (a similar pattern as hippocampus but with reduced amplitude), and caudate evokes co-fluctuations involving control and salience/ventral attention systems. These observations posit a dynamic explanation for the observed specificity of static cortico-subcortical functional connections [19, 49].

### Movies decouple sensorimotor systems from one another

We extended our study of system-level events from resting-state to scans where subjects passively watched clips from movies in the scanner–i.e. naturalistic stimuli [50, 51]. In line with [29], we found that system-level events tended to occur during the resting periods between successive movie scenes. We also found that, compared to rest, the timing of events in sensorimotor and attentional networks tended to decouple from one another during movie-watching, untethering brain systems from one another and allowing them to flexibly, and with greater autonomy, track and process audiovisual stimuli. This observation explains the reduced amplitude of system events during movie-watching compared to rest within those systems during their respective system events. Peak co-fluctuations for sensorimotor/attention networks occur at similar times during rest, effectively amplifying the mean RMS at that instant. During moviewatching, however, the decoupling disrupts the coincidence of events so that fewer systems reach their peak coincidentally, thereby reducing the brain-wide RMS amplitude during those systems’ events.

Interestingly, this decoupling phenomenon is not exhibited uniformly throughout the brain. While most edges exhibit reductions in their co-fluctuation magnitude (MOVIE-REST) a small subset exhibit increases. Cortically, these edges are concentrated in higher-order association systems, e.g. default mode and control networks [25]. However, the strongest and most consistent increases were centered on cerebellar sub-regions (lobule X; vermal and hemispheric parts [26]) and their links to sensorimotor and attention networks and the caudate. This is of particular interest, as the FC of lobule X to cerebral cortex has been largely uncharted and/or no strong connections reported [52, 53], but may be linked to movement and motion [54]. This hypothesis is broadly in line with recent studies suggesting that the cerebellum may play an active role in mental simulation of action, e.g. while viewing complex naturalistic stimuli, repurposing cortical areas to support this function [55, 56].

### “A poor man’s” edge communities

A growing number of studies have applied data-driven clustering algorithms to static FC to partition brain territories into communities, modules, and systems [57, 58]. The result is a “hard” partition of neural elements into non-overlapping systems (but see [59, 60] for notable examples). Perhaps the key advantage of hard partitions is their simplicity: a node is or is not affiliated with a given system and the boundaries between partitions are unambiguous. The structure of the underlying network data, however, is not unambiguous and is characterized by heterogeneity at the level of nodes’ connectivity profiles, which generally extend beyond the boundaries of their respective systems. Moreover, because the clusters defined by hard partitions have inherited functionally evocative names-e.g. visual, somatomotor, attention systems-they have inadvertently reinforced a view of nervous system organization in which modules (and by extension, their constituent brain regions) are associated with one function and one function only.

Inspired by applications in network science [61, 62], several recent studies have proposed “edge-centric” approaches for defining and characterizing the brain’s system-level architecture [30, 63–65]. Unlike traditional methods for defining brain network systems, the “edge-centric” approach assigns cluster labels at the level of edges so that, from the perspective of each node, it maintains an affiliation to a plurality of clusters. This type of overlapping organization-in which there exists few well-defined clusters-has been termed “pervasive overlap” and is a hallmark of many real-world systems [66]. It also more closely agrees with established neuroscience, where cells and populations respond to a range of stimuli (even in areas traditionally viewed as uni-functional).

However, one of the challenges associated with the obtaining edge-level cluster assignments is that it has a large computational burden, requiring the user to generate and partition high-dimensional edge time series or edge functional connectivity [30, 63] (but see [67] for an alternative approach). The method developed here circumvents the need to create memory intense data structures and the implementation of computationally costly clustering algorithms, aligning each edge with the system whose event evokes the greatest co-fluctuation magnitude (on average). Although we use pre-defined system labels here, this approach is generally agnostic to their origins; it would be reasonable to align edge time series with systems estimated at the subject level [18] or using data-driven clustering algorithms, e.g. infomap or modularity maximization.

Note that our approach for establishing edge-level cluster assignments requires that we first identify system-level events, which involves estimating a null distribution of RMS values for each system that, depending on the number of systems and resolution of parcellation, can still be computationally costly in a large multi-subject dataset. Fortunately, edge system assignments could be estimated easily using other methods. For instance, one could calculate the correlation of individual edges time series with system-level edge time series, assigning edges to the system with the maximum correlation magnitude (Fig. S16).

### Events as a continuum *versus* a discrete category

The focus of previous analyses of edge time series has been, largely, on the highest amplitude events defined globally. This choice of analysis is justified because, all things equal, higher-amplitude frames *must* contribute more to the time-averaged pattern of FC; FC is calculated as the average of an edge time series, so frames with greater amplitude are necessarily weighted more heavily. However, several recent studies have noted that sub-event frames–those that would not traditionally be categorized as an event–contain useful information about an individual [12, 28] and, with the exception of those with very low amplitudes (which are more likely to be contaminated by motion [17]), are also predictive of static FC [68]. These findings suggest that the dichotomy of “events” *versus* “non-events” likely is an oversimplification.

Here, we shift focus away from the whole-brain and onto individual systems. Although we still study the behavior of these systems through an event-based framework for reasons noted above, by emphasizing individual systems, we can interrogate smaller systems whose behavior might be washed out and overwhelmed by that of larger systems at the global scale. To this end, our approach and results are broadly in line with the view that a strict discretization of frames into “event” and “nonevent” categories overlooks an underlying and largely continuous distribution of co-fluctuation amplitude. A potentially fruitful direction for future research is to examine not only system levels events, but the entire spectrum of co-fluctuation amplitudes from the perspective of brain systems.

### Future directions

The results of this study and the system-based framework for examining edge time series presents several avenues for future research. Here, we examined edge time series and system-level events during rest and movie-watching and, because this is the first study to do so, focused largely at the group level. Future studies should investigate system-level effects during other conditions, e.g. during tasks, and in at the individual level in different populations, e.g. developmental and/or clinical.

Here, we generated representative co-fluctuation maps for each system as the average over all events originating in that system. While these maps offered a summary of the central tendency for each edge’s co-fluctuation magnitude, it does not capture the heterogeneity across samples. In previous studies, we showed that global events could be clustered into a small number of repeating patterns or “states” [11, 17, 28]. It is unclear from the results presented here whether similar cluster structure exists at the level of systems. Future studies should investigate this in greater detail.

Another possible direction for future work involves detecting different categories of events. Here, we identify events at a system level by isolating moments in time when the edges within canonical brain systems exhibit stronger than expected co-fluctuations. However, we could also consider other categories of events, including points in time with off-diagonal blocks–i.e. between-system connections–exhibit stronger than expected co-fluctuations. This might be especially useful in experimental conditions wherein we expect systems to couple with one another to perform cognitively demanding tasks [69].

A final and potentially fruitful direction for future research is to move beyond correlation measures–two systems tend to exhibit events simultaneously–and examine their causal relationships with respect to one another. For instance, is there evidence of “gating” mechanisms, such that given an event in system *s* at time *t*, a global event (or an event in system *s*′ ≠ *s*) is likely to follow.

### Limitations

A massive literature has been assembled documenting time-varying changes in spontaneous fMRI BOLD activity and connectivity [70] along with its behavioral [71] and clinical importance [72]. These findings are complemented by a growing number of studies at other spatiotemporal scales using invasive imaging techniques to reveal meaningful time-varying fluctuations in micro- and meso-scale spontaneous activity [73–75]. Despite this preponderance of evidence, several recent studies have suggested that some of the apparent dynamism of fMRI BOLD activity is, in fact, not-dynamic at all and directly anticipated by the static correlation structure–i.e. FC [68, 76, 77]. In general, these studies estimate an empirical FC matrix and use it to simulate synthetic and random multivariate fMRI BOLD data. When analyzed, these time series exhibit dynamic properties similar to those observed in the original data, leading the authors to conclude that analogous features in the observed data reflect random fluctuations around a ground truth static correlation structure.

Although the results of these studies help constrain hypotheses about time-varying networks, they also have limitations. First, if time-varying network features, including events, were truly stochastic fluctuations, then their intersubject correlation should approach a floor value. That is, the timing of events between two subjects should be uncorrelated. However, the results presented here and in our previous studies [9, 10, 29] indicate that both global RMS and events synchronize during movie-watching. These observations indicate that exogenous stimuli play a role in organizing the temporal structure of high-amplitude brain activity. Second, these studies raise a deeper philosophical issue concerning the origins of brain activity and how best to model it. It is generally accepted that spontaneous brain activity is generated by an anatomically constrained dynamical system [16, 78, 79]. The correlation structure of this activity–i.e. FC–is a convenient summary statistic and useful marker, but is fundamentally epiphenomenal and plays no causal role in generating future patterns of brain activity (at least over short timescales). In other words, null models in which data are generated so as to preserve a specific pattern are of correlations are not only inconsistent with the biophysical underpinnings of fMRI BOLD data, but possibly circular (assuming that the observed correlation structure is the generator of itself).

In short, the question of whether spontaneous fMRI BOLD data exhibits non-stationarities or meaningful time-varying fluctuations in connectivity remains contentious and unresolved [80]. Here, we adopt the view that there exists neurobiologically meaningful information in time-varying co-fluctuation patterns.

A second potential limitation concerns the decision to analyze fMRI BOLD data processed without global signal regression (GSR). Following other recent studies [22, 81], we chose to exclude GSR from the processing steps (while carefully monitoring and correcting for in-scanner motion and other artifacts), as GSR can artifactually induce anti-correlations [82, 83] and because the global signal contains information about brain-wide, population level activity [84] and arousal [85].

A final concern is the role that the temporal signal to noise ratio (tSNR) in each system might impact the detectability of events, the rate at which they appear, and their amplitude. Here, we did not explicitly quantify its role, but note that inferior cortical systems, e.g. limbic, and subcortical areas have lower tSNR in the HCP dataset. Future studies should investigate how tSNR, along with system size and static properties of FC, impact events.

## MATERIALS AND METHODS

### Datasets, preprocessing, and time series extraction

The Human Connectome Project (HCP) dataset [21] consists of structural magnetic resonance imaging (T1w), resting state functional magnetic resonance imaging (rsfMRI) data, and movie watching functional magnetic resonance imaging (mwfMRI) from young adult subjects. The dataset spans two collection sites, at Washington University in St. Louis on a 3T MRI machine and at the Center for Magnetic Resonance Research at the University of Minnesota on a 7T MRI machine. For the present study, the 3T data consist of a subset of 100 unrelated adults (“100 Unrelated Subjects” published by the Human Connectome Project) and the 7T data consist of 172 adults (including twins and siblings). The study was approved by the Washington University Institutional Review Board and informed consent was obtained from all subjects. HCP 3T and 7T data were quality controlled based on motion summary statistics and visual inspection. After exclusion of four high motion subjects and one subject due to software error, the final HCP 3T subset consisted of 95 subjects the final subset utilized included 95 subjects (56% female, mean age = 29.29 ± 3.66, age range = 22-36). After motion exclusion criteria, the final HCP 7T subset consisted of 129 subjects (60% female mean age = 29.36 ±3.36, age range = 22-36; 75 unique families).

Data from the Midnight Scan Club was used as a confirmation analysis and had also been collected with parameters detailed in [23]. Data included 10 healthy, young adult participants (50% females; age range = 24-34) and were recruited at Washington University. Informed consent was obtained from each participant and the study was approved by the Washington University School of Medicine Human Studies Committee and Institutional Review Board. A Siemens TRIO 3T MRI scanner was used over 12 separate runs at midnight and collected over multiple days. Ultimately, 5 hours of resting state fMRI data were collected for each participant.

A comprehensive description of the HCP imaging parameters and image prepocessing can be found in [86] and in HCP’s online documentation (https://www.humanconnectome.org/study/hcp-young-adult/document/1200-subjects-data-release). For all HCP subjects (even those in the 7T dataset), T1w were collected on a 3T Siemens Connectome Skyra scanner with a 32-channel head coil. Subjects underwent two T1-weighted structural scans, which were averaged for each subject (TR = 2400 ms, TE = 2.14 ms, flip angle = 8°, 0.7 mm isotropic voxel resolution). For all fMRI data collected, 4 gradient-echo planar imaging sequences were collected for each subject: two runs were acquired with posterior-to-anterior phase encoding direction and two runs were acquired with anterior-to-posterior phase encoding direction.

HCP 3T fMRI was collected on a 3T Siemens Connectome Skyra with a 32-channel head coil. Each resting state run duration was 14:33 min, with eyes open and instructions to fixate on a cross (TR = 720 ms, TE = 33.1 ms, flip angle = 52°, 2 mm isotropic voxel resolution, multiband factor = 8). HCP 7T fMRI was collected on a 7T Siemens Magnetom scanner with a 32-channel head coil. Four movie watching data runs were collected, each lasting approximately 15 minutes (frames = 921, 918, 915, 901), with subjects passively viewing visual and audio presentations of movie scenes (TR = 1000 ms, TE = 22.2 ms, flip angle = 45^°^, 1.6 mm isotropic voxel resolution, multi-band factor = 5, image acceleration factor = 2, partial Fourier sample = 7/8, echo spacing = 0.64 ms, bandwidth = 1924 Hz/Px). Movies consisted of both freely available independent films covered by Creative Commons licensing and Hollywood movies prepared for analysis [87].

Structural and functional HCP images were minimally preprocessed according to the description provided in [86]. 7T data were downloaded after correction and reprocessing announced by the HCP consortium in April, 2018. Briefly, T1w images were aligned to MNI space before undergoing FreeSurfer’s (version 5.3) cortical reconstruction workflow. fMRI images were corrected for gradient distortion, susceptibility distortion, and motion, and then aligned to the corresponding T1w with one spline interpolation step. This volume was further corrected for intensity bias and normalized to a mean of 10000. This volume was then projected to the 2mm *32k_fs_LR* mesh, excluding outliers, and aligned to a common space using a multi-modal surface registration [88]. The resultant CIFTI file for each HCP subject used in this study followed the file naming pattern: `*_Atlas_MSMAll_hp2000_clean.dtseries.nii`.

Structural and functional MSC images were minimally preprocessed using fMRIPrep version 20.2.0. Briefly, this pipeline includes intensity correction, skull stripping, volumetric segmentation and spatial normalization of T1w images. Functional data were slice time, motion, and “fieldmap-less” distortion corrected, and aligned to T1w images. More complete descriptions of fMRIPrep’s pipeline can be found at [89] or in fMRIPrep’s online documentation (https://fmriprep.org/en/20.2.0/citing.html). The pipeline was run using the following options: FreeSurfer (version 6.0.1); “fieldmap-less” susceptibility distortion correction; the NKI skull stripping template; slice-time correction; six degrees of freedom for the BOLD to T1w co-registration. Functional outputs of the pipeline were captured as CIFTI files.

All resting state and moving watching fMRI images were nuisance regressed in the same manner. Each minimally preprocessed fMRI was linearly detrended, band-pass filtered (0.008-0.25 Hz), confound regressed and standardized using Nilearn’s `signal.clean` function, which removes confounds orthogonally to the temporal filters. Multiple confound regression strategies were employed for various analyses of the current manuscript. The 36-parameter confound regression strategy included six motion estimates, mean signal from a white matter, cerebrospinal fluid, and whole brain mask, derivatives of these previous nine regressors, and squares of these 18 terms; this strategy is one that includes GSR. The aCompCor strategy included six motion estimates, derivatives of these previous six regressors, and squares of these 12 terms, in addition to five anatomical CompCor components [90]; aCompCor is a non-GSR strategy. For HCP 3T, data from both strategies were used at different points of the manuscript. For MSC data and for both HCP 7T, rest and movie fMRI were regressed the aCompCor strategy. Following these preprocessing operations, the mean signal was taken at each time frame for each node, forming nodal time series. Cortical nodes were defined by the Schaefer 200 cortical parcellation [25] in the *32k_fs_LR* surface space. Volumetric ROIs of subcortical [27] and cerebellar [26] regions were defined in the MNI152 2mm space, conforming to the CIFTI specification.

## EDGE TIME SERIES

We used a combination of cortical, cerebellar, and subcortical parcellations to divide brains into *N* = 482 regions of interest. FC between regions *i* and *j* is operationalized as a correlation coefficient, so that:

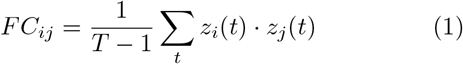

where **z**_*i*_ = [*z_i_*(1),…, *z_i_*(*T*)] is the vector of z-scored activity recorded from region *i*.

The time series for edge {*i*, *j*} is calculated by simply omitting the summation and normalization step. That is, *e_ij_*(*t*) = *z_i_*(*t*) · *z_j_*(*t*). Intuitively, *e_ij_*(*t*) indicates the magnitude of co-fluctuation between regions *i* and *j* at time *t*. Its sign is either positive or negative depending on whether those regions are simultaneously deflecting above or below their respective means. Its magnitude is large if both regions exhibit big deflections and small if the deflections are similarly close to zero.

Importantly, edge time series (ETS) are a precise decomposition of FC into its framewise contributions. The sum ∑_*t*_ *e_ij_*(*t*) is proportional to the static FC between regions *i* and *j*; the proportionality is made exact if normalized by *T* – 1.

Here, we modify the procedure for calculating ETS in a subtle, yet important, way. Normally, we would calculate the z-scored activity of region *i* at time *t* as 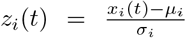, where 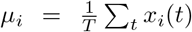 and 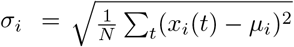. In short, both the sample mean and standard deviation entail averaging over all frames. However, the value of activity, *x_i_*(*t*) can vary with in-scanner movement, impacting our estimates of the mean and standard deviation. Accordingly, we omit any frame flagged for high motion when we estimate *μ_i_* and *σ_i_*. This means that the “z-scored” time series generally have means and variances that are not precisely 0 and 1, respectively. In principle, frame censoring would be a preferred approach for addressing issues related to motion. However, the null model used for event detection requires that we not drop any frames, motivating the procedure described above.

### Event detection

Previous studies have shown that whole-brain cofluctuation patterns averaged over a small sub-set of high-amplitude frames can explain a large fraction of variance in static FC. Following those studies, we detect these “events” by comparing co-fluctuation magnitudes against a null model. Events are defined by first identifying contiguous blocks of supra-threshold frames (whose magnitude is statistically greater than that of the null model) and identifying, within those frames, the one with the greatest amplitude.

In greater detail, we calculate amplitude as the square root of the mean squared co-fluctuations (RMS). To detect global events, we calculate RMS as:

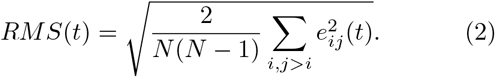

We also calculated a system level analog of this measure. For system, s, we calculate its local RMS as:

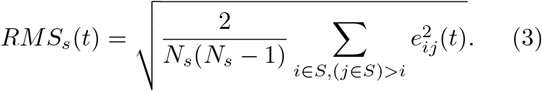

Here, *S* is the set of all regions assigned to system *s* and *N_s_* = |*S*| is the size of said system.

To detect events, we compared these observed RMS values against a null distribution obtained by randomly circularly shifting each region’s activity time series and calculating, from the shifted data, the ETS and RMS values. Note that this null model exactly preserves the mean and variance of regions’ activity while approximately preserving its spectral properties. However, it destroys correlation structure.

For each frame, *t*, we calculated the proportion of the null distribution that exceeded the value *RMS*(*t*). This proportion served as a non-parametric *p*-value. We corrected for multiple comparisons by adjusting the critical value so as to preserve a false discovery rate of *q* = 0.01. For each contiguous block of frames in which all RMS values exceeded that of the null, we assessed whether any of those frames had been flagged for high levels of motion. If so, we omitted the entire block from subsequent analysis. For blocks with no flagged frames, we identified the frame with the largest *RMS* and treated this as the representative co-fluctuation pattern for that block. All analyses were carried out on these peak frames.

### Overlapping system labels

For each system, *s*, we can identify frames, *τ*, at which *RMS_s_*(*τ*) is statistically greater than that of null model and greater than all nearby supra-threshold frames. We consider these frames to be system events, in that they occur when a particular system – rather than the brain as a whole – exhibits a significant increase its in co-fluctuation amplitude.

Although these events are detected based on the behavior of an individual system, *s*, we can still examine the whole-brain co-fluctuation pattern that is expressed at these instants. Specifically, we can aggregate the co-fluctuation magnitude for all node pairs, *i* and *j*, at time *t* = *τ*. These patterns can be averaged over all occurrences to obtain a mean co-fluctuation pattern that characterizes the behavior of the entire brain at the time that system s has an event. Doing so for all systems yields a [*N* × *N* × *N_sys_*] arrays.

One of the interesting features of this matrix is that we can collapse across its third dimension (systems) to identify for each node pair, {*i*, *j*}, the system whose event elicits the strongest co-fluctuation. The index of this system serves as an edge-level system label. Note that alternative procedures have been described elsewhere for obtaining edge clusters, but those methods require either constructing an unwieldy *edge* × *edge* matrix and using data-driven methods to cluster edges [30], or using similar methods to cluster edge time series directly [63].

### Regional entropy

Based on the edge-level system assignments, we can calculate for each node the extent to which its edges deviate from uniformity. Specifically, we can calculate that region’s entropy. To do so, we first calculate *p_is_*: the fraction of edges incident upon *i* assigned to system *s*. We can then calculate that region’s entropy as:

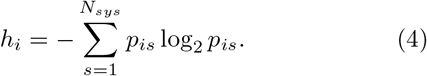

If the distribution of edge system assignments are concentrated within a small subset of systems, then *h_i_* → 0. As the distribution of assignments approaches uniformity, *h_i_* increases. We normalize *h_i_*, bounding it to the interval [0, 1] by dividing by log_2_(*N_sys_*).

### Subject-specific system labels

Throughout the main text we focus on “canonical brain systems” and areal labels defined *a priori* based on previous studies [25–27]. Although these system boundaries and labels offer a generally agreed-upon nomenclature for referencing functional systems and have reasonable out-of-sample validity, we also wanted explore the extent to which these labels could be made subject-specific.

To this end, we performed the following analysis. For each participant, we first aggregated their four restingstate scans to obtain a single representative FC matrix. Next, for each subject we calculated a group-averaged FC matrix (after excluding, from the aggregated data, that subject’s matrix), and calculated, based on that group matrix, the mean connectivity fingerprint for each system. Then, using that subject’s own FC matrix, we identified for each node the system-level fingerprint to which its own fingerprint was most similar, assigning nodes to the corresponding system.

### Data-driven detection of modules

We also used a data-driven approach to detect group-level modules. First, we estimated a group-level FC matrix by averaging subject-specific matrices. Next, we used multi-scale modularity maximization to detect communities of different sizes and compositions. Modularity maximization [91, 92] is a popular technique for community detection in which communities are defined as groups of nodes whose density of connections to one another maximally exceeds what would be expected by chance. This intuition is formalized by the modularity quality function, *Q*, defined as:

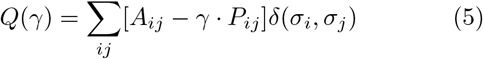

where *A_ij_* and *P_ij_* are the observed and expected weights of the connection between nodes *i* and *j*, *γ* is a resolution parameter that scales the relative importance of *P_ij_* with respect to *A_ij_*, and *σ_i_* is the community assignment of node *i*. The function *δ*(*σ_i_*, *σ_j_*) is the Kronecker delta function and is equal to 1 when *σ_i_* = *σ_j_* and 0 otherwise. The aim of modularity maximization is to select nodes’ community assignments – the values of *σ_i_* for all *i* – such that *Q*(*γ*) is as big as possible.

Here, we sampled 100000 values of *γ* from the distribution of all connection weights greater than —0.05 in the group-averaged FC matrix. For each sample, we added a small amount of random noise (between ±1% of its original value). We then used a custom implementation of the Louvain algorithm to optimize *Q*(*γ*). This procedure resulted in 100000 independent estimates of communities at different topological scales.

We grouped *γ* values into 15 percentile-based non-overlapping bins. In each bin, we calculated the module co-assignment matrix, whose element *D_ij_* denotes the fraction of all partitions assigned to that bin in which nodes *i* and *j* were assigned to the same community. We then used a consensus clustering algorithm to define a single representative partition for each bin based on its corresponding co-assignment matrix. Briefly, the algorithm worked by clustering the co-assignment matrix, assessing whether the new set of clusters are identical and, if not, calculating a new co-assignment matrix and repeating this procedure. The clustering algorithm used here was, again, modularity maximization. However, we adapted the problem to co-assignment matrices, so that, in the modularity equation, *A_ij_* and *P_ij_* represented the observed and expected co-assignment probabilities (the value of *P_ij_* was calculated as the expected co-assignment had we observed the same set of partitions but with nodes randomly assigned to communities). This procedure resulted in 15 consensus partitions representing different topological scales. We excluded the first and last consensus partitions as they corresponded to a partition in which most nodes were assigned to a single large community and their own communities, respectively.

---

**FIG. S1.**
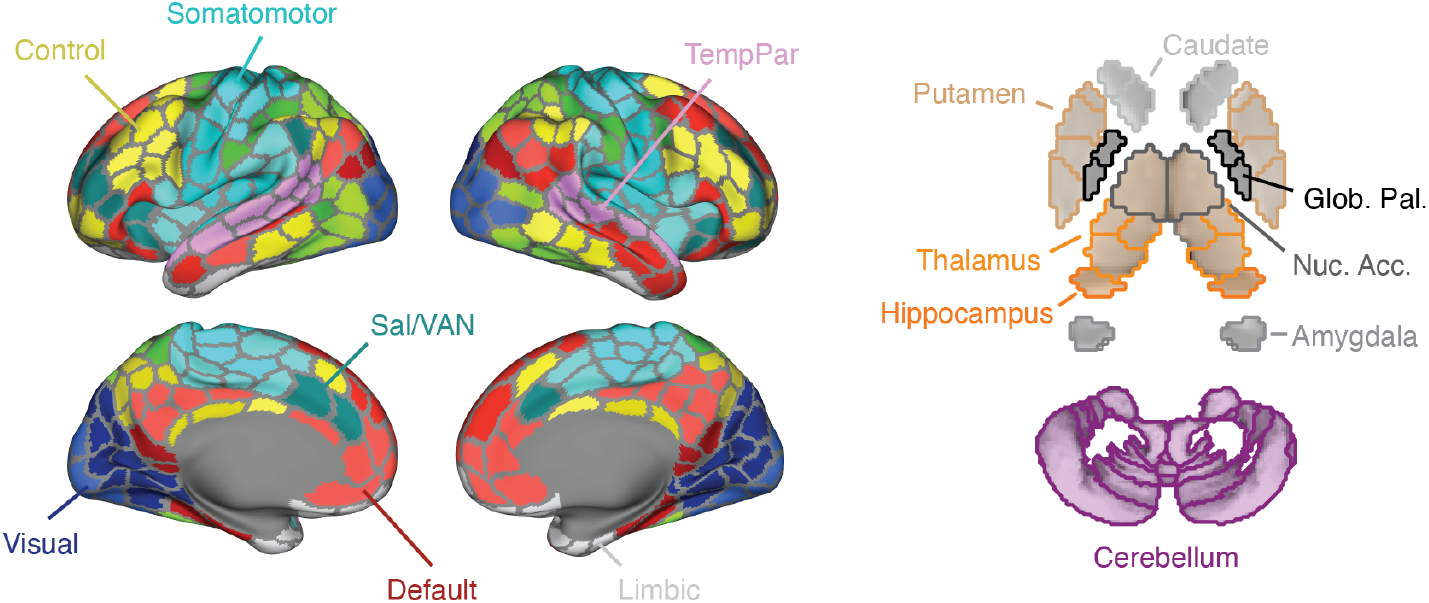
System and areal levels. Cortical systems from [25], cerebellar divisions from [26], and subcortical divisions from [27].

**FIG. S2.**
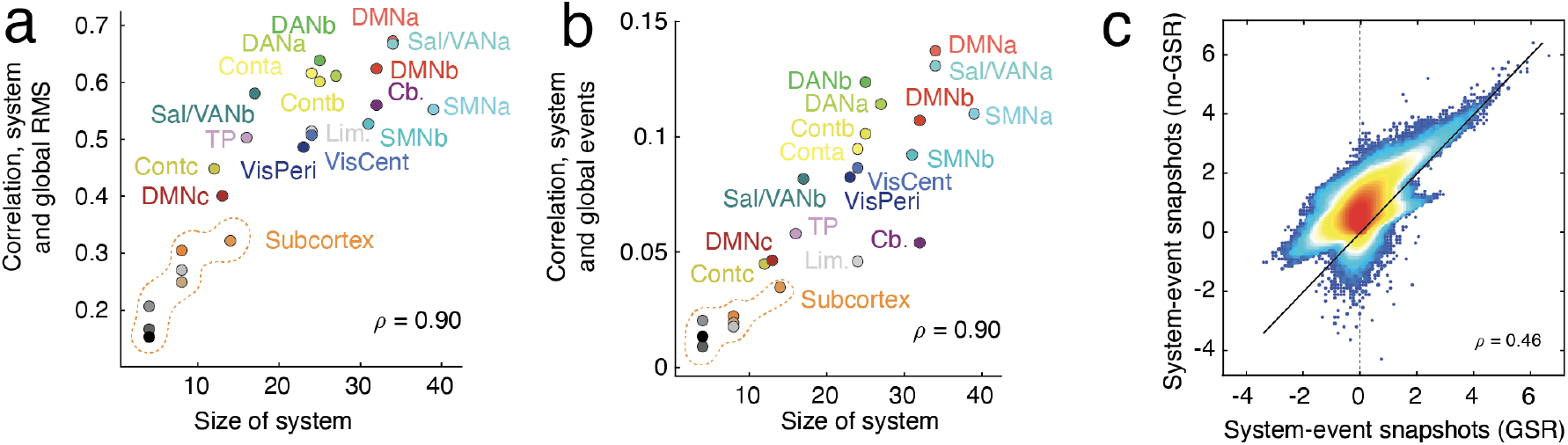
Comparison between system event co-fluctuation patterns with and without global signal regression. (*a*) Population-averaged correlation of system RMS with global RMS *versus* system size (number of regions). (*b*) Population-averaged correlation of system event timing with global events *versus* system size (number of regions). (*c*) We pooled the elements of system event co-fluctuation matrices across all systems for fMRI data processed with and without global signal regression. We then plotted these elements against one another and found a broad correspondence.

**FIG. S3.**
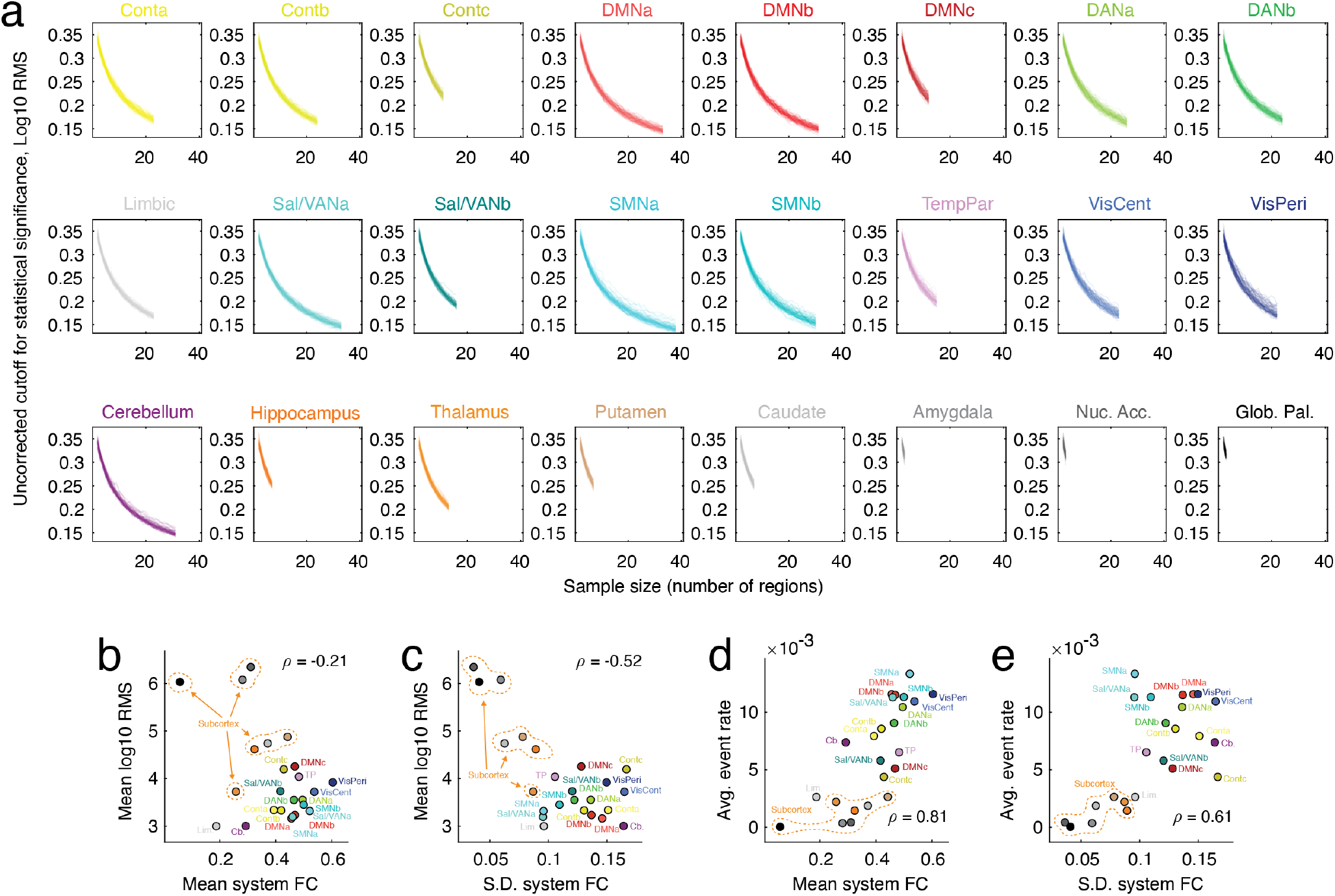
System RMS as a function of system size. For each system, we calculated its mean RMS after sub-sampling different numbers of nodes (*a*) Plots for all cortical, cerebellar, and sub-cortical systems. Relationship of RMS and event rate with properties related to FC. (*b*) Correlation between RMS and mean within-system static FC. (*c*) Correlation between RMS and standard deviation of within-system static FC. (*d*) Correlation between event rate and mean within-system static FC. (*e*) Correlation between event rate and standard deviation of within-system static FC.

**FIG. S4.**
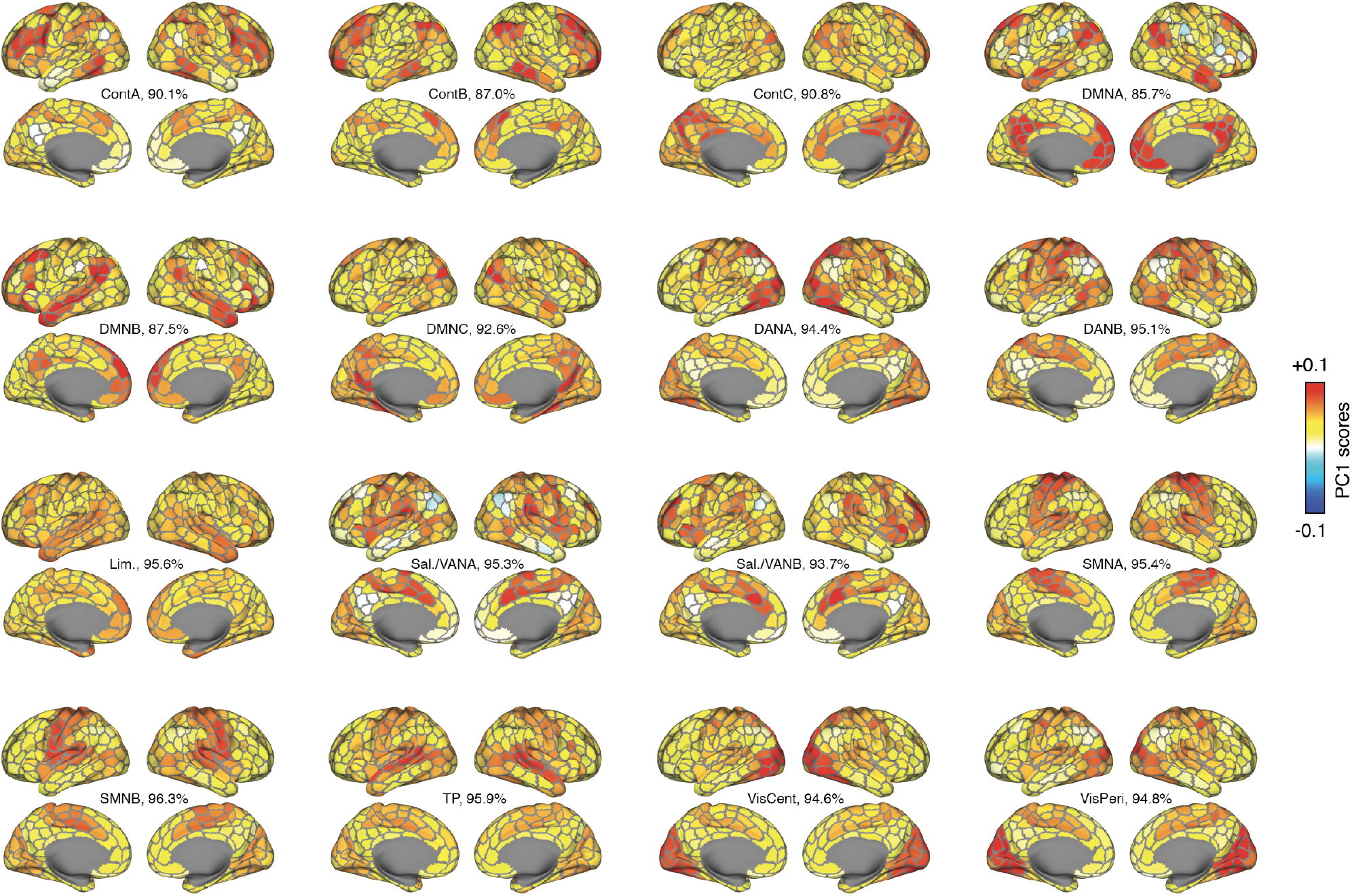
Projections of system event co-fluctuation matrices onto brain regions. For each cortical system, we used singular value decomposition to decompose the group-averaged system event co-fluctuation matrix. Here, we show the first principal component for each system projected onto the cortical surface along with the total variance explained by that component.

**FIG. S5.**
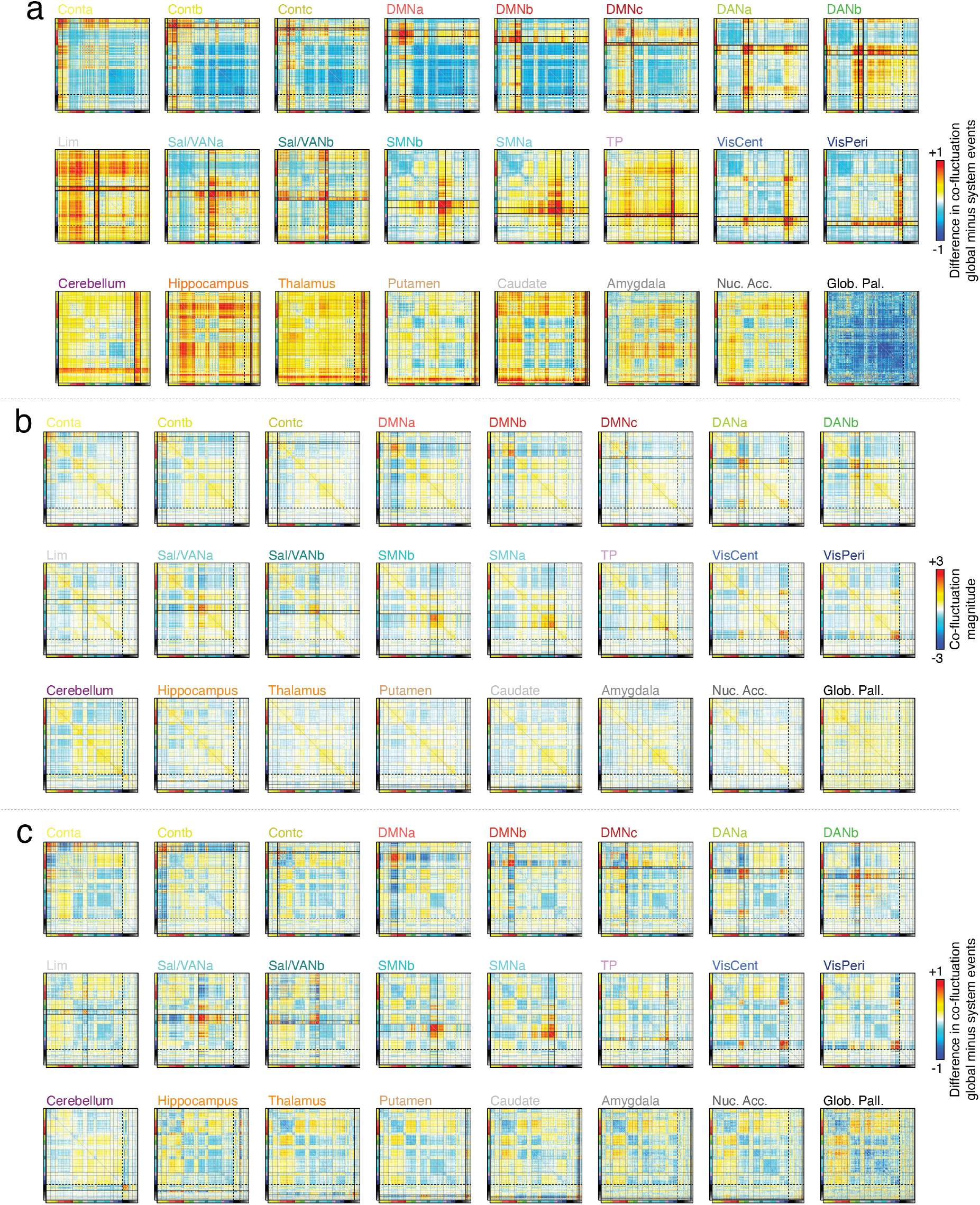
System event co-fluctuation maps. (*a*) Difference in system event co-fluctuation patterns and global co-fluctuation for fMRI data processed without global signal regression (same data described in the main text). (*b*) System event co-fluctuation matrix for fMRI data processed with global signal regression. (*c*) Difference in system event co-fluctuation patterns and global co-fluctuation for fMRI data processed with global signal regression.

**FIG. S6.**
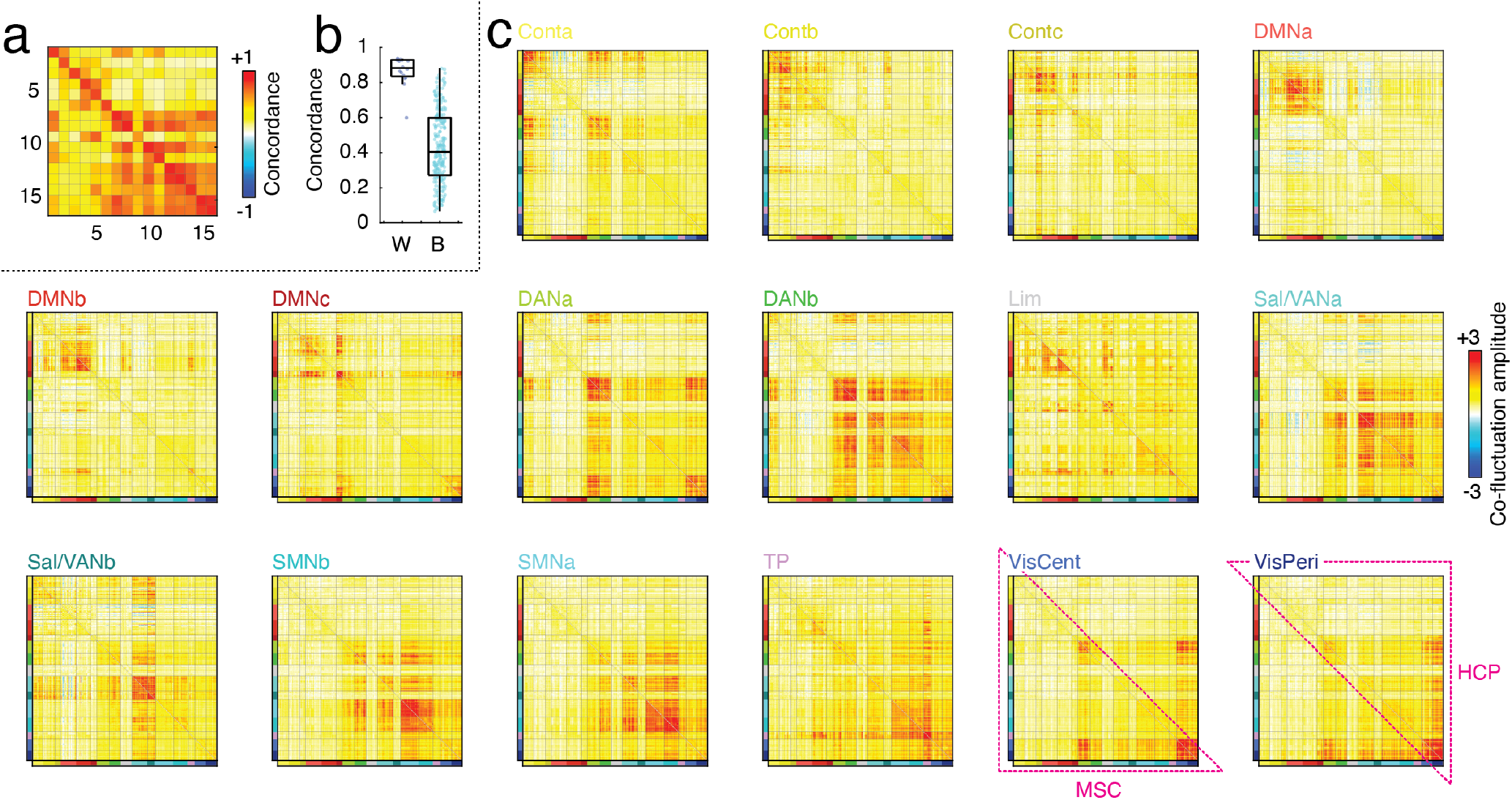
Comparing system event snapshots between HCP and MSC datasets. We calculated system event co-fluctuation patterns for the HCP dataset (main text) and the midnight scan club (MSC). (*a*) The similarity matrix of co-fluctuation patterns between HCP (rows) and MSC (columns) datasets. (*b*) The same-system similarity (diagonal) exhibited significantly greater level of similarity than the between-system elements. (*c*) Example co-fluctuation matrices. The upper triangle shows co-fluctuation values from the HCP dataset; the lower triangle depicts MSC co-fluctuation values.

**FIG. S7.**
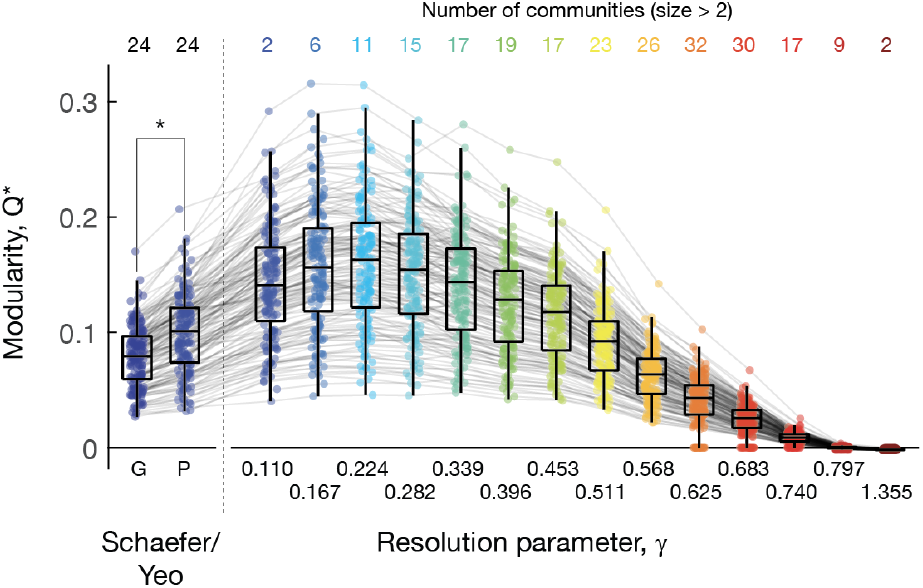
Modularity of static FC for different system-level partitions. In the main text we calculated system-level events based on a “canonical” set of systems from [25]. These systems were identical for all subjects. We also estimated a version of these systems that were fit to each subjects’ own FC pattern as well as a series of multi-scale, data-driven estimates. Here, we compare these system labels in terms of their ability to partition subjects’ static FC into segregated subsystems by calculating the modularity, *Q*\* induced by the partition [93].

**FIG. S8.**
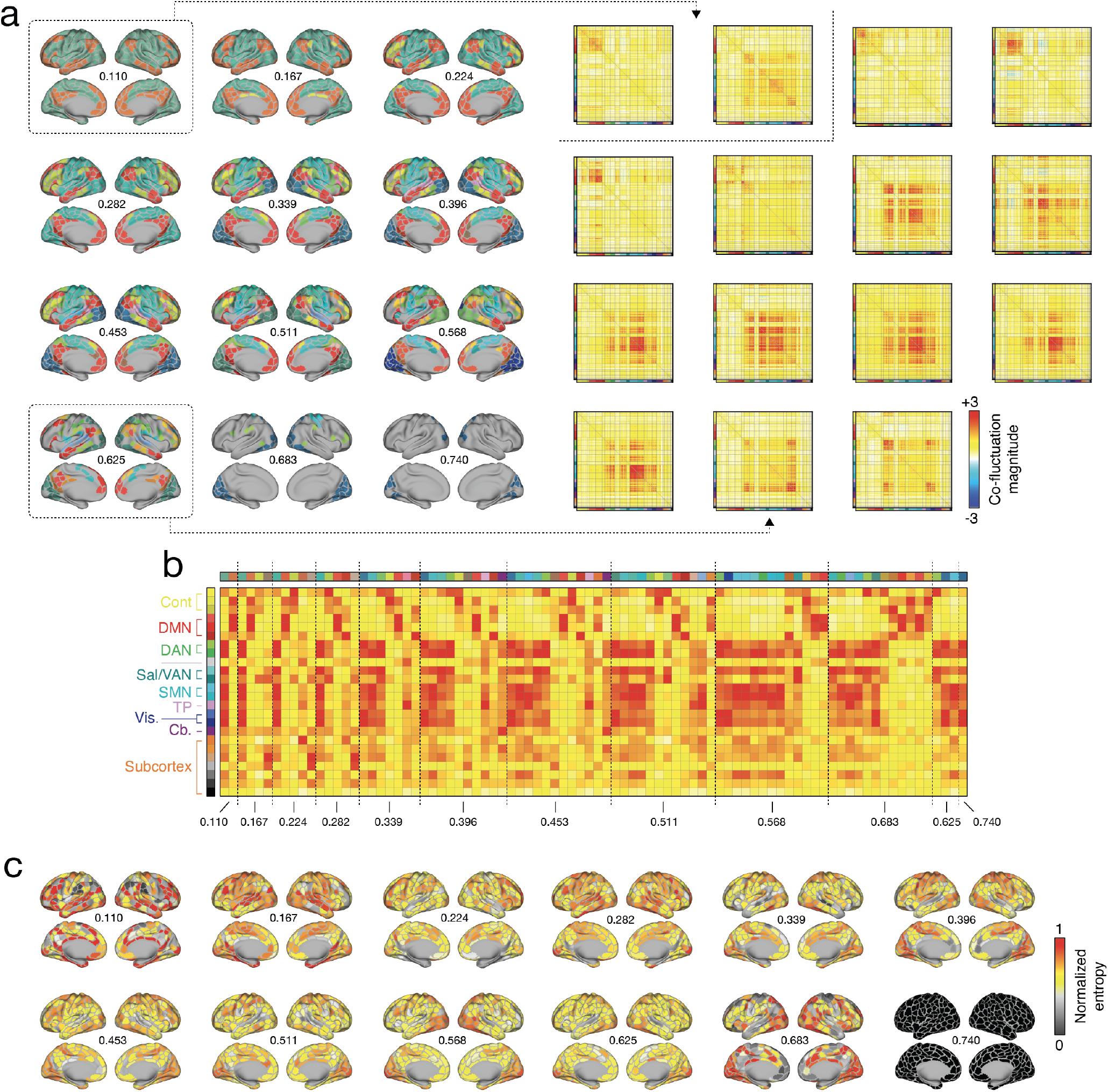
Data-driven community summary. (*a*) Detected communities at different hierarchical levels (*left*) and example co-fluctuation matrices for two resolutions (*γ* ≈ 0.110 and *γ* = 0.625). (*b*) Similarity of system event co-fluctuation patterns reported in main text with analogous patterns estimated for detected communities. (*c*) Induced.

**FIG. S9.**
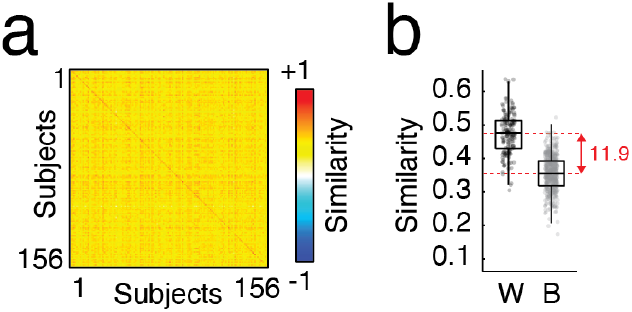
Subject-specificity of system-level co-fluctuation patterns. (*a*) Mean similarity between co-flucutation magnitudes estimated based on data from scans REST1 and REST2. (*b*) Diagonal elements of the matrix *versus* off-diagonal.

**FIG. S10.**
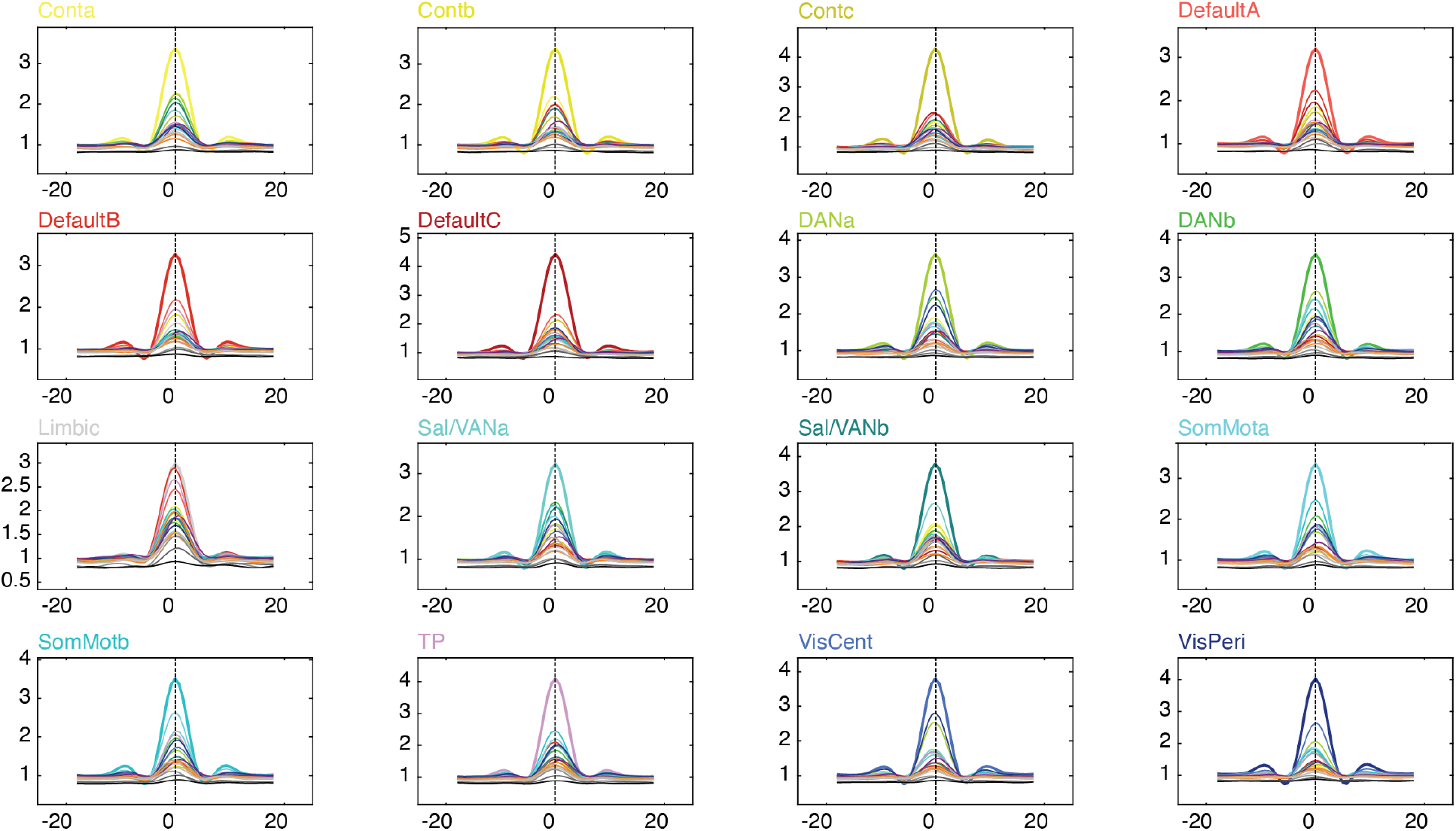
Mean event timing. RMS curves for system events. Event time is set equal to *t* = 0.

**FIG. S11.**
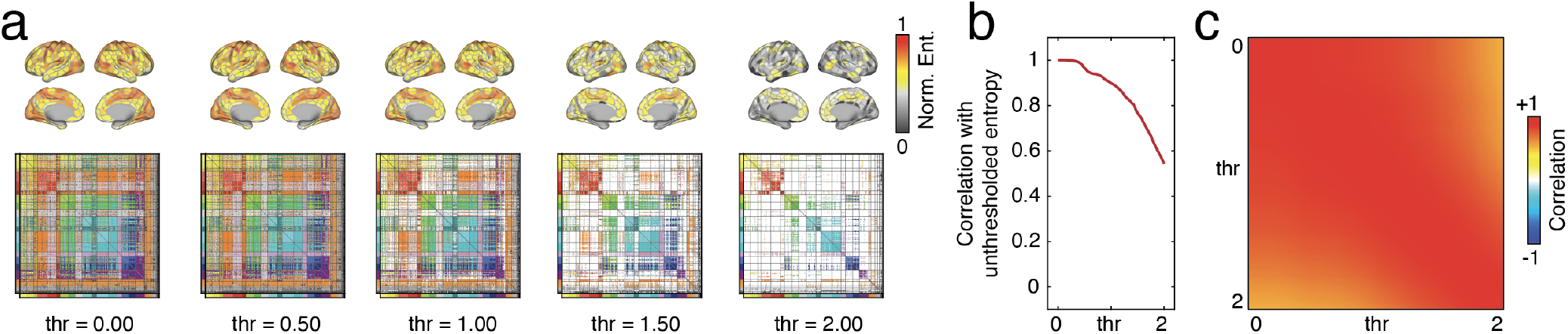
Effect of sparsity on overlap patterns. (*a*) Edge community matrix (*bottom*) thresholded to preserve only the labels of edges whose maximum co-fluctuation exceeds that of a threshold. Here, we show thresholds from 0 to 2 in increments of 0.5. The top panel shows the overlap pattern (entropy) induced by the thresholded communities. (*b*) Spatial correlation of the entropy map at different thresholds with respect to unthresholded map. (*c*) Pairwise similarity of entropy maps at thresholds from 0 to 2.

**FIG. S12.**
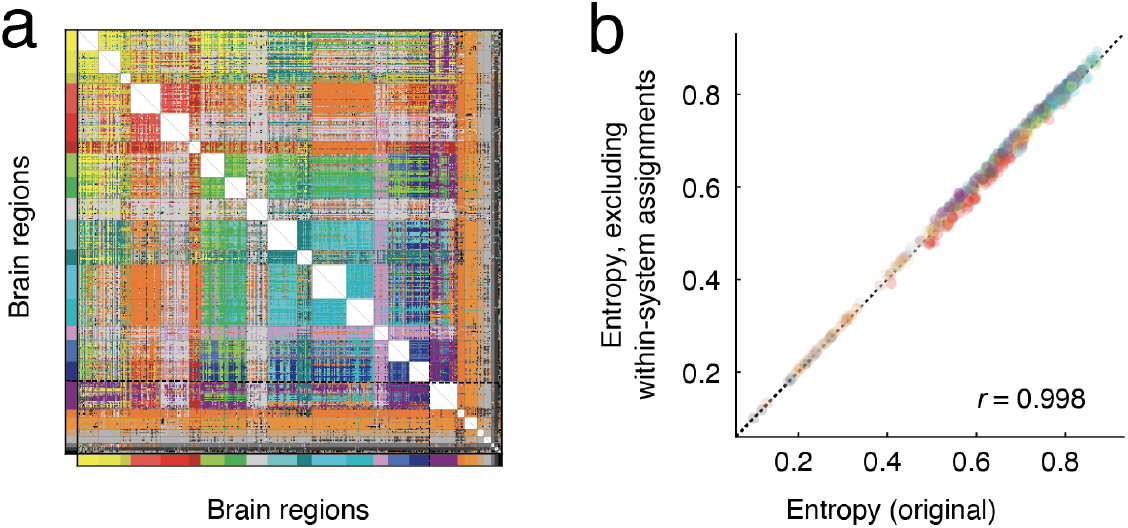
Edge cluster assignments and entropy after excluding within-system edges. We removed and ignored all within-system edges and recalculated regional entropy. We found an extremely close correspondence with the entropy values reported in the main text, suggesting that within-system edges (that might be trivially associated with their respective system) do not drive the overlap.

**FIG. S13.**
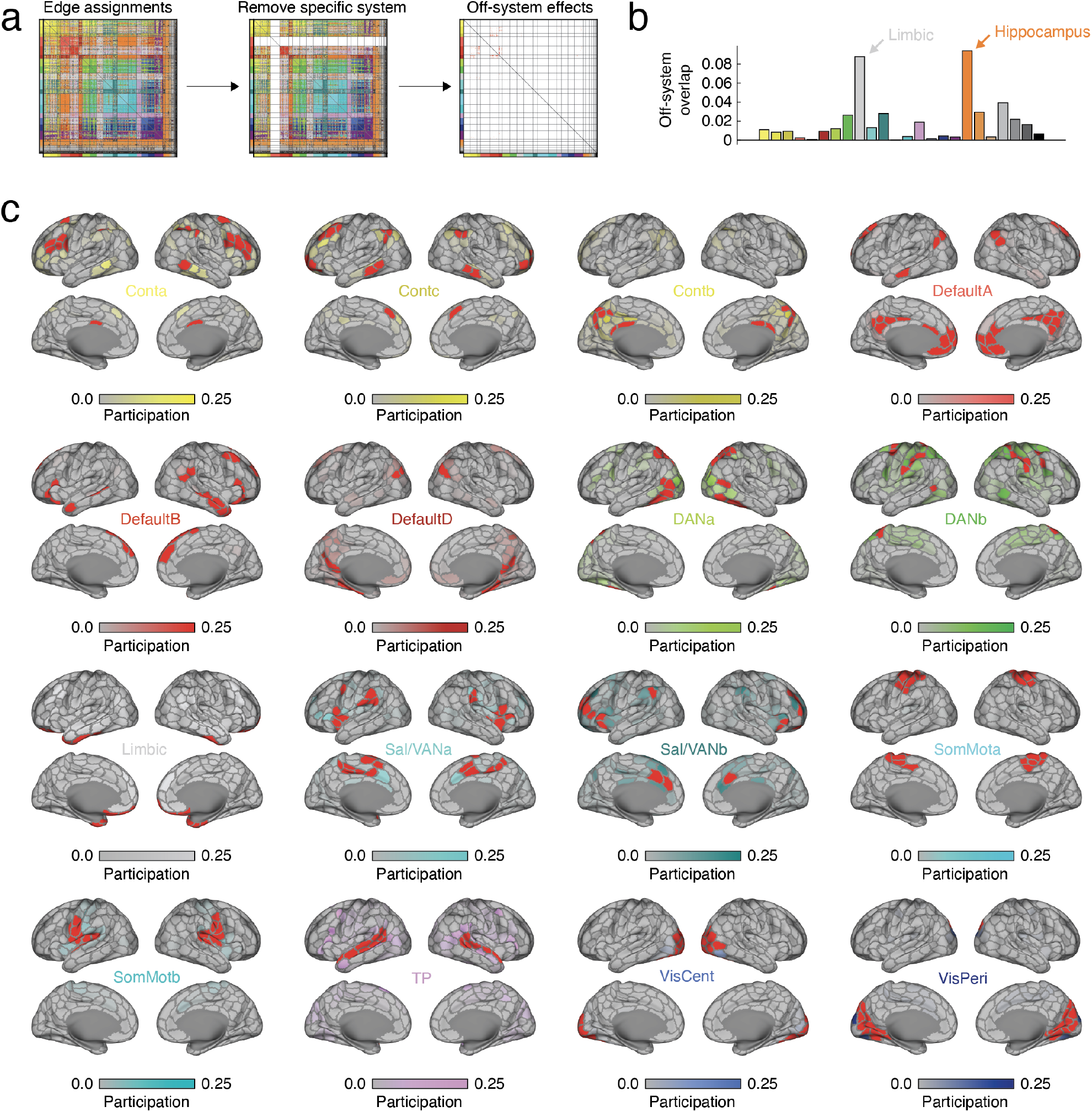
Cortical affiliation after removing connections incident upon each system. (*a*) Edge community matrix before (*left*) and after (*middle*) removing all edges incident upon a specific system (in this case, DMNa). Once edges are removed, we calculated each node’s affiliation with the system (*right*). (*b*). Mean overlap across brain systems. (*c*) Overlap pattern projected onto cerebral cortex. Red regions indicate those whose corresponding rows/columns were removed (and artificially set to 0).

**FIG. S14.**
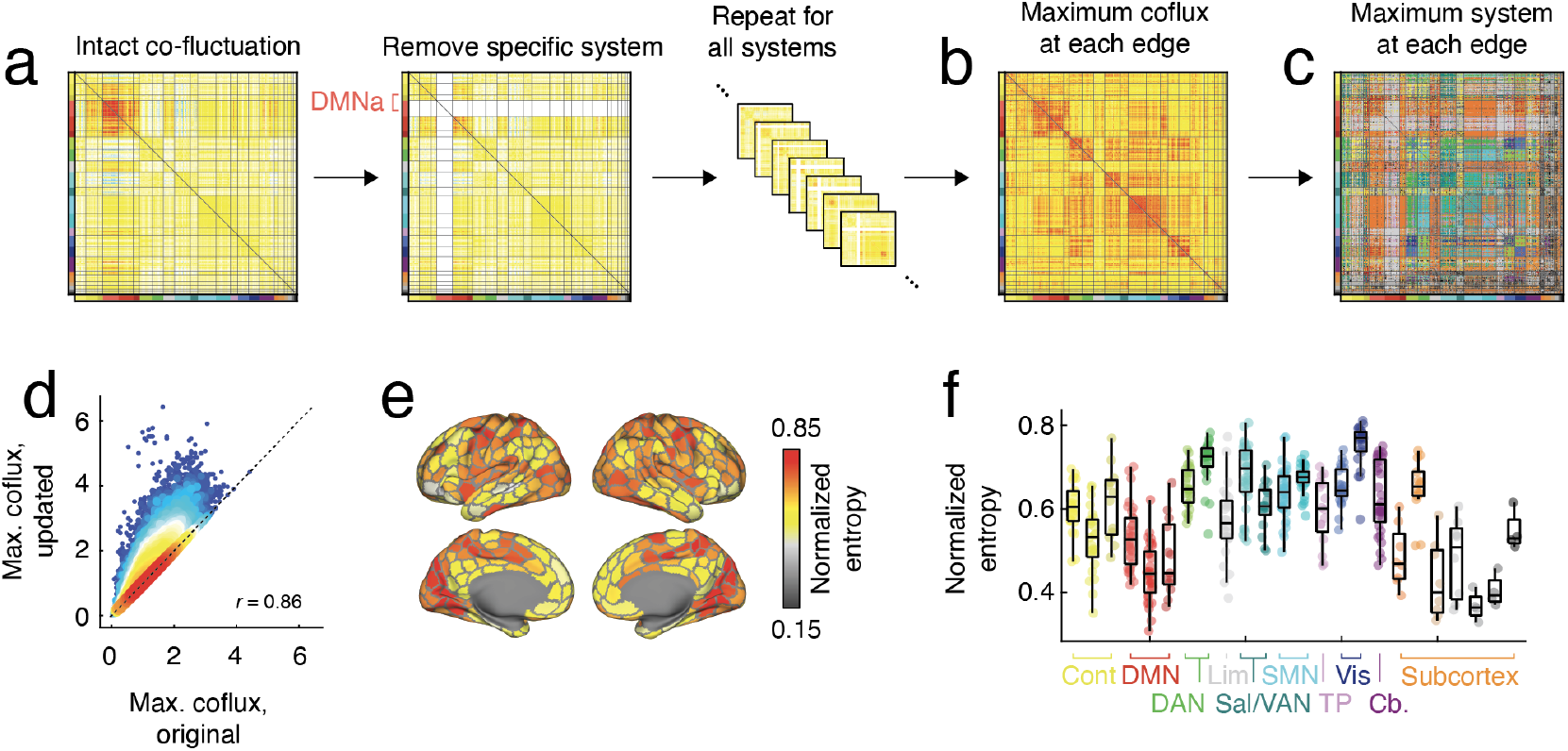
Cortical affiliation after removing connections to incident upon said system. (*a*) After excluding edges incident upon each system, we calculated, based on the remaining systems (*b*) the maximum co-fluctuation and (*c*) the corresponding edge community label. (*d*) We compared the maximum co-fluctuation after removing systems with the maximum co-fluctuation based on the intact (unthresholded) matrix. (*e*) Entropy maps projected onto cortical surface. (*f*) Entropy maps aggregated by brain systems.

**FIG. S15.**
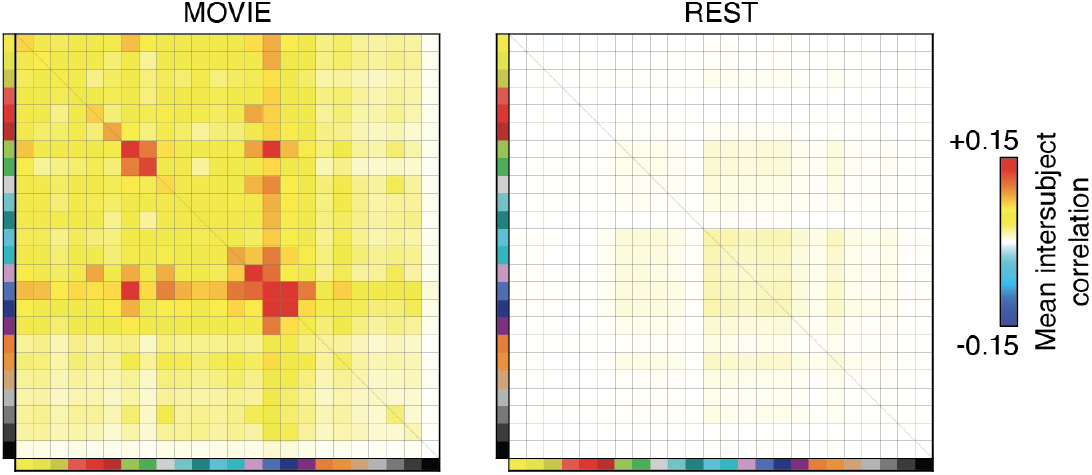
Inter-subject and inter-system correlations during rest and movie-watching. We calculated system-level RMS curves for movie-watching and resting-state scans. (*a*) Mean inter-subject correlation matrix (averaged over all unique pairs of subjects) for movies and (*b*) rest.

**FIG. S16.**
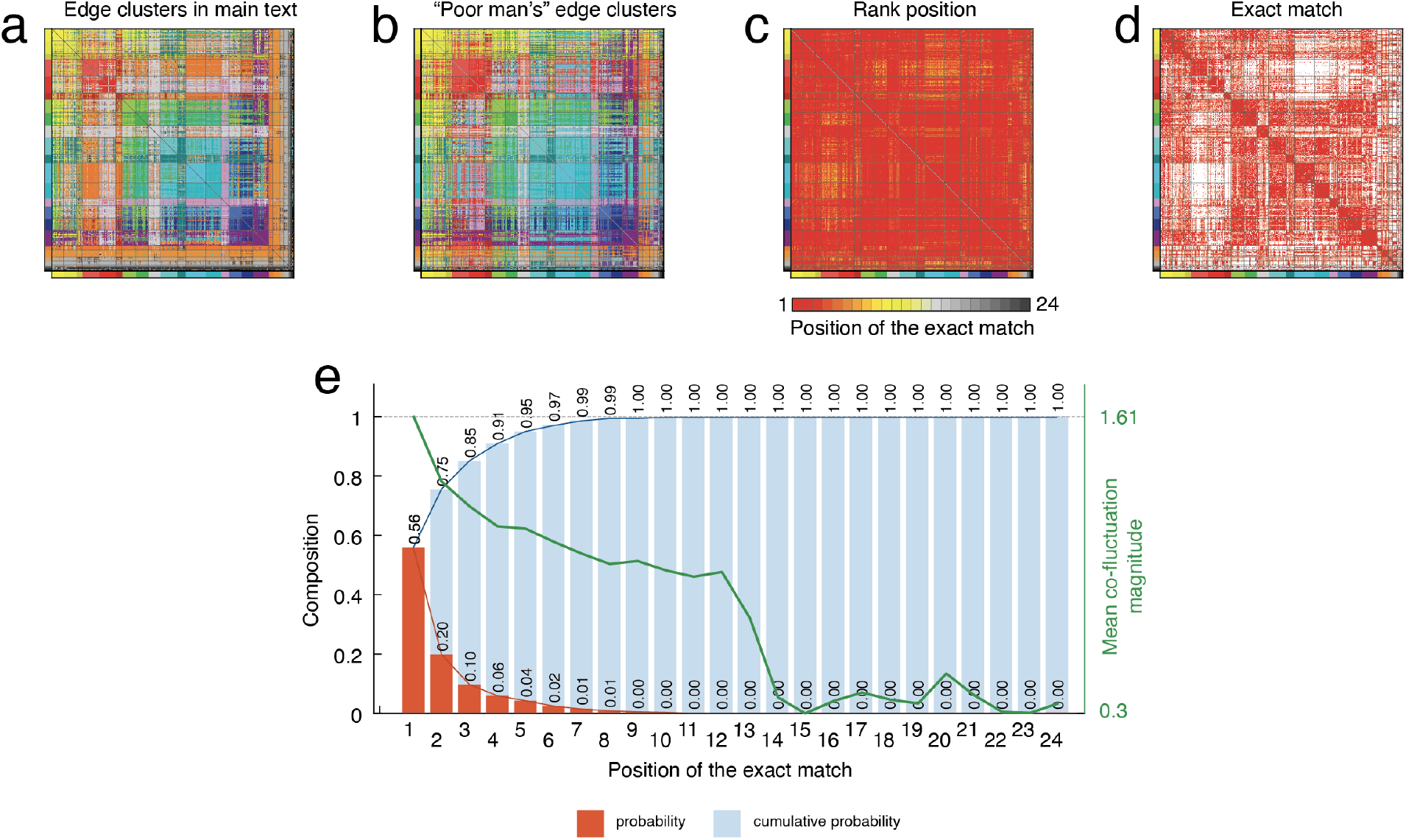
Alternative method for estimating edge-level system assignments. In the main text, we assigned edges to brain systems based on their relative magnitude during “system event snapshots”. Here, we present an alternative method that does not necessitate the detection of events, which requires estimating a null distribution for RMS values for each system. Briefly, in this approach we calculate the mean time course for edges within each brain system, yielding 24 time courses. Next, we identify which of these 24 time courses each edges’ time series is maximally correlated and assign that edge to that system. (*a*) Edge clusters reported in the main text. (*b*) Edge clusters estimated using the alternative method. (*c*) Position (rank) of the edge clusters in the main text among the 24 systems that each edge is correlated against. A value of “1” indicates that the edge clusters match exactly-i.e. the strongest correlation is the system with the strongest co-fluctuation amplitude during its event; a value of “24” indicates a maximal mismatch. (*d*) Exact matches-i.e. rank of “1”. (*e*) Frequency and cumulative distribution of rank values. Overlaid (in green) is the mean co-fluctuation amplitude of edges at each rank.

